# Individual Variation in Control Network Topography Supports Executive Function in Youth

**DOI:** 10.1101/694489

**Authors:** Zaixu Cui, Hongming Li, Cedric H. Xia, Bart Larsen, Azeez Adebimpe, Graham L. Baum, Matt Cieslak, Raquel E. Gur, Ruben C. Gur, Tyler M. Moore, Desmond J. Oathes, Aaron Alexander-Bloch, Armin Raznahan, David R. Roalf, Russell T. Shinohara, Daniel H. Wolf, Christos Davatzikos, Danielle S. Bassett, Damien A. Fair, Yong Fan, Theodore D. Satterthwaite

## Abstract

The spatial distribution of large-scale functional networks on the anatomic cortex differs between individuals, and is particularly variable in networks responsible for executive function. However, it remains unknown how this functional topography evolves in development and supports cognition. Capitalizing upon advances in machine learning and a large sample of youth (n=693, ages 8-23y) imaged with 27 minutes of high-quality fMRI data, we delineate how functional topography evolves during youth. We found that the functional topography of association networks is refined with age, allowing accurate prediction of an unseen individual’s brain maturity. Furthermore, the cortical representation of executive networks predicts individual differences in executive function. Finally, variability of functional topography is associated with fundamental properties of brain organization including evolutionary expansion, cortical myelination, and cerebral blood flow. Our results emphasize the importance of considering both the plasticity and diversity of functional neuroanatomy during development, and suggest advances in personalized therapeutics.

## INTRODUCTION

During childhood, adolescence, and young adulthood, the human brain must develop to support increasingly complex cognitive and behavioral capabilities. One broad domain of cognition that undergoes particularly protracted development is executive function, which encompasses diverse cognitive processes including working memory, performance monitoring, and task switching (Best and Miller, 2010; Gur et al., 2012). Individual differences in executive function have been linked to meaningful functional outcomes such as academic achievement (Arffa, 2007; Best et al., 2011), and deficits of executive function are associated with violence, initiation of drug use, and risk taking behaviors (Reynolds et al., 2019). Executive dysfunction is also associated with most major neuropsychiatric diseases (Shanmugan et al., 2016), including attention deficit hyperactivity disorder and psychosis (Barkley, 1997; Wolf et al., 2015).

Executive processes rely upon a spatially distributed set of brain regions that span frontal, parietal, and temporal cortex (Alvarez and Emory, 2006; Niendam et al., 2012; Rottschy et al., 2012). These regions have low cortical myelin content (Glasser and Van Essen, 2011), receive a disproportionate amount of cerebral blood flow (Satterthwaite et al., 2014b; Taki et al., 2011), and have greater areal expansion compared to other cortical regions in humans (Reardon et al., 2018) and analogous regions in non-human primates (Hill et al., 2010). Non-invasive studies using functional MRI (fMRI) in humans have shown that these distributed regions activate together during cognitively demanding executive tasks and also show coherent signal fluctuations at rest (Cole and Schneider, 2007; Marek and Dosenbach, 2018; Satterthwaite et al., 2013b), allowing them to be understood as large-scale functional networks. Typically, these networks have been compared across individuals by alignment with brain structure, which assumes that there is a stable correspondence between functional and structural anatomy across individuals (Laumann et al., 2015). However, recent evidence from multiple independent efforts has demonstrated that there is marked inter-individual variability in the spatial topography of functional brain networks even after accurate alignment of brain structure (Bijsterbosch et al., 2018; Braga and Buckner, 2017; Glasser et al., 2016; Gordon et al., 2017a; Gordon et al., 2017b; Gordon et al., 2017c; Kong et al., 2018; Laumann et al., 2015; Li et al., 2019; Wang et al., 2015).

Studies of highly-sampled individuals for whom numerous sessions of scanning data were acquired have established that an individual’s functional topography is highly reproducible across scanning sessions (Gordon et al., 2017c; Laumann et al., 2015). Furthermore, several studies have reported that topographic variability across individuals is maximal in brain networks responsible for executive functioning (Gordon et al., 2017b; Gordon et al., 2017c; Kong et al., 2018; Li et al., 2019; Wang et al., 2015). This finding aligns with work showing that these same association networks also show the greatest inter-individual variation in their connectivity profiles (Gratton et al., 2018; Kong et al., 2018; Li et al., 2019; Mueller et al., 2013), and can be used for accurate identification of individuals (Finn et al., 2015; Miranda-Dominguez et al., 2014). Understanding subject-specific functional topography also allows prediction of an individual’s spatial pattern of activation across diverse tasks (Gordon et al., 2017c; Laumann et al., 2015; Li et al., 2019; Tavor et al., 2016; Wang et al., 2015). Failure to account for such individual variation in functional topography may lead differences in spatial distribution to be aliased into measurement of inter-regional functional connectivity, potentially biasing both inference and interpretation (Bijsterbosch et al., 2018; Li et al., 2019).

Despite such rapidly accruing evidence for the importance of individual differences in functional neuroanatomy, to our knowledge no studies have characterized variation of functional topography in youth. To address this gap, here we tested three inter-related hypotheses. First, we hypothesized that functional topography would be systematically refined during development, with developmental changes being concentrated in association cortex. Second, we predicted that variation in the functional topography of executive networks would predict individual differences of executive functioning. Third and finally, we anticipated that these developmental changes and associations with executive functioning would be constrained by fundamental properties of brain organization, including evolutionary expansion and cortical myelination. To test these hypotheses, we capitalized upon recent advances in machine learning and a large sample of youth who participated in the Philadelphia Neurodevelopmental Cohort (PNC; Satterthwaite et al. (2014a)).

## RESULTS

We studied 693 youths aged 8-23 years who completed imaging as part of the PNC (**Supplementary Figure 1**) with over 27 minutes of high-quality fMRI data (see Supplementary Methods). To delineate person-specific functional networks, we used a spatially-regularized form of non-negative matrix factorization (NMF; Lee and Seung (1999)) that has previously been shown to accurately identify functional networks in individuals (Li et al., 2017). This approach involved three steps (**Figure 1**). In the first step, a group atlas was created by running NMF on the concatenated timeseries of a sub-sample of 100 subjects. In the second step, to ensure reproducibility, the group atlas was re-created on a total of 50 sub-samples (n = 100 participants each), and a consensus set of networks was derived using spectral clustering. In the third step, individualized networks were identified for each participant by iteratively applying NMF to each participant’s data, with the consensus networks used as a prior to ensure correspondence across participants.

**Figure 1.**
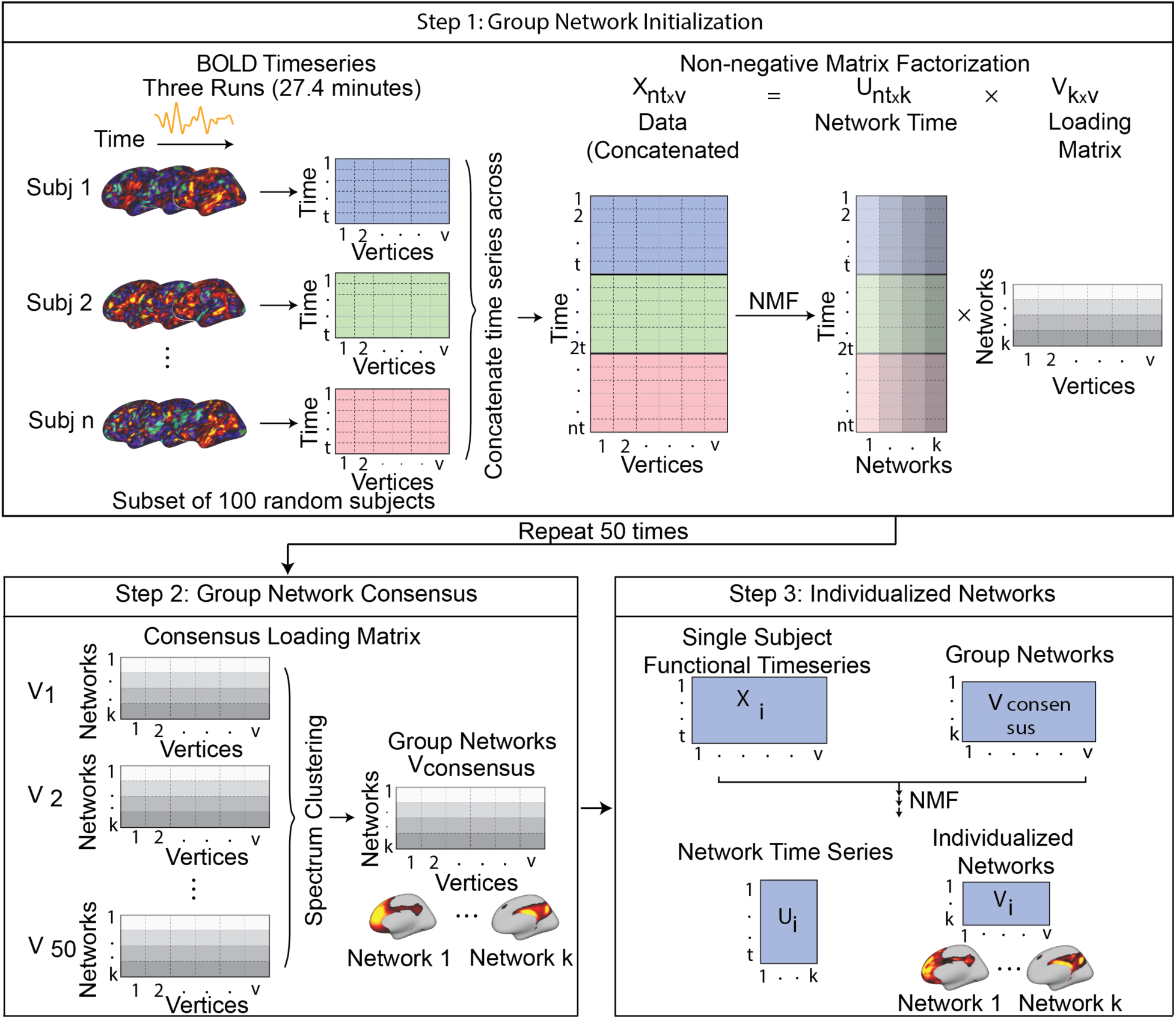
Schematic of spatially regularized non-negative matrix factorization (NMF) for individualized network parcellation. Each subject had three fMRI runs; we concatenated these for each subject, resulting in a 27.4 minutes timeseries with 555 time points for each subject. In the first step, we randomly selected 100 subjects and concatenated their time series into a matrix with 55,500 time points (rows) and 18,715 vertices (columns). Non-negative matrix factorization was used to decompose this matrix into a timeseries matrix and loading matrix. The loading matrix had 17 rows and 18,715 columns, which encoded the membership of each vertex at each network. This procedure was repeated 50 times, with each run including a different subset of 100 subjects. In the second step, a normalized cut-based spectral clustering method was applied to cluster the 50 loading matrices into one consensus loading matrix, which served as the group atlas and ensured correspondence across individuals. In the third step, NMF was used to calculate individualized networks for each participant, with the group atlas used as a prior.

To facilitate comparison to other methods (Kong et al., 2018), we identified 17 functional networks in each participant (**Figure 2**). In contrast to methods that discretely assign each vertex to a single network, NMF yields a probabilistic (soft) parcellation. This probabilistic parcellation can be converted into discrete (hard) network definitions for both display and comparison with other methods by labeling each vertex according to its highest loading. Visual inspection suggested that these discretized networks showed a high correspondence with a widely-used 17-networks solution (Yeo et al., 2011). To quantitatively compare these atlases, as well as subsequent analyses of individual parcellations and other cortical properties (see below), we used a spatial permutation test that relies on random surface-based rotations (or “spins”) to test the significance of spatial correlation between brain maps (Alexander-Bloch et al., 2018; Gordon et al., 2016). This conservative statistical procedure preserves the spatial covariance structure of the data and provides a more appropriate null distribution than randomly shuffling surface locations (see Supplementary Methods). Using this approach, we found significant alignment (*P_spin_* < 0.001) between our group atlas and the canonical 17 networks (Yeo et al., 2011).

**Figure 2.**
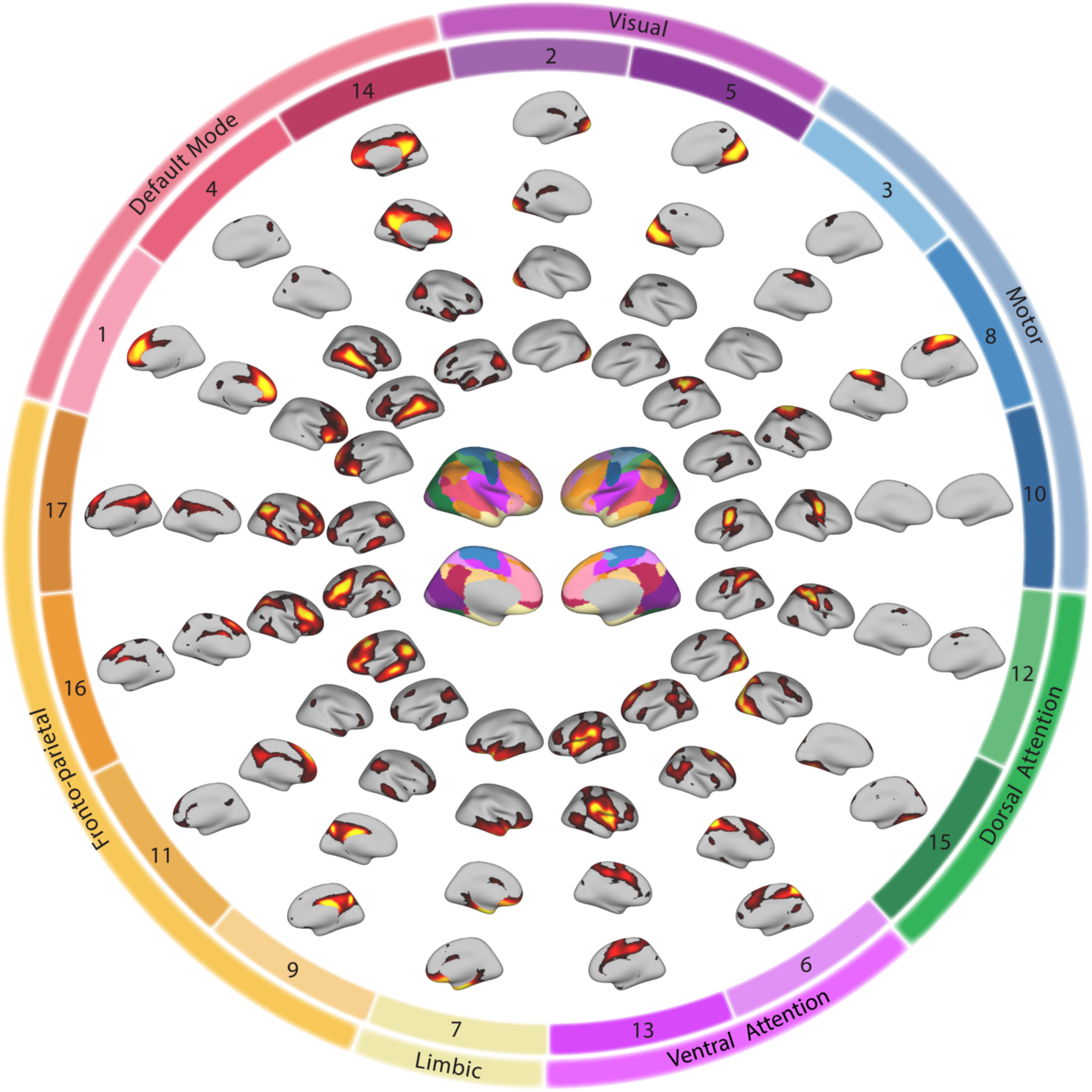
Group atlas of 17 networks. Our NMF-based method for network parcellation relies on a group atlas as a prior for identifying networks in individuals. In this atlas, there are 17 loadings for each vertex, which quantify the extent it belongs to each network. The networks in the group atlas include medial and lateral visual networks (numbers 2 and 5); hand, foot and face motor networks (numbers 3, 8, and 10); dorsal attention networks (number 12 and 15), ventral attention networks (numbers 6 and 13); a limbic network (number 7); fronto-parietal control networks (numbers 9, 11, 16, and 17), and default mode networks (numbers 1, 4, and 14). For display, vertices can be assigned to the network with the highest loading, yielding a discrete network parcellation (center).

### Individualized networks improves functional homogeneity compared to group averaged networks

Next, this group atlas was tailored to each individual’s data using NMF, providing subject-specific networks. As in previous work (Gordon et al., 2016; Kong et al., 2018), we evaluated the quality of these individualized networks by calculating the homogeneity of the functional timeseries within each network. The mean within-network homogeneity for individualized networks using NMF was significantly higher than in randomly rotated networks (*P_spin_* < 0.001). Furthermore, homogeneity within NMF-based individualized networks was higher than that in either the NMF-based group atlas or the standard 17-network group atlas (**Supplementary Figure 2**). As an additional validation step, we compared our NMF-based method with a recently-introduced method which uses a multi-session hierarchical Bayesian model (MS-HBM; Kong et al. (2018)) to identify individualized networks. The mean homogeneity of MS-HBM in our sample was numerically lower than our NMF based method, and nearly identical to a prior application of MS-HBM to adults (0.31 vs. approximately 0.32; Kong et al. (2018)). Furthermore, we found that the individual parcellations provided by NMF and MS-HBM were significantly aligned (*P_spin_* < 0.001). These initial results suggest that single-subject parcellations provide an improved fit to each participant’s data compared to standard atlases that do not consider variation in functional neuroanatomy.

### Across-subject variability of network topography is maximal association cortex

Visual examination of many individual subjects revealed that while the gross spatial distribution of networks was consistent across participants, distinct person-specific topographic features could be readily observed (**Figure 3**). Consistent with prior reports (Gordon et al., 2017b; Gordon et al., 2017c; Kong et al., 2018; Li et al., 2019; Mueller et al., 2013; Wang et al., 2015), heterogeneity in the spatial distribution of networks was particularly apparent in association networks such as the fronto-parietal control, ventral attention, and default mode networks. In contrast, participant-level representations of somatomotor and visual networks appeared to be much more consistent across individuals.

**Figure 3.**
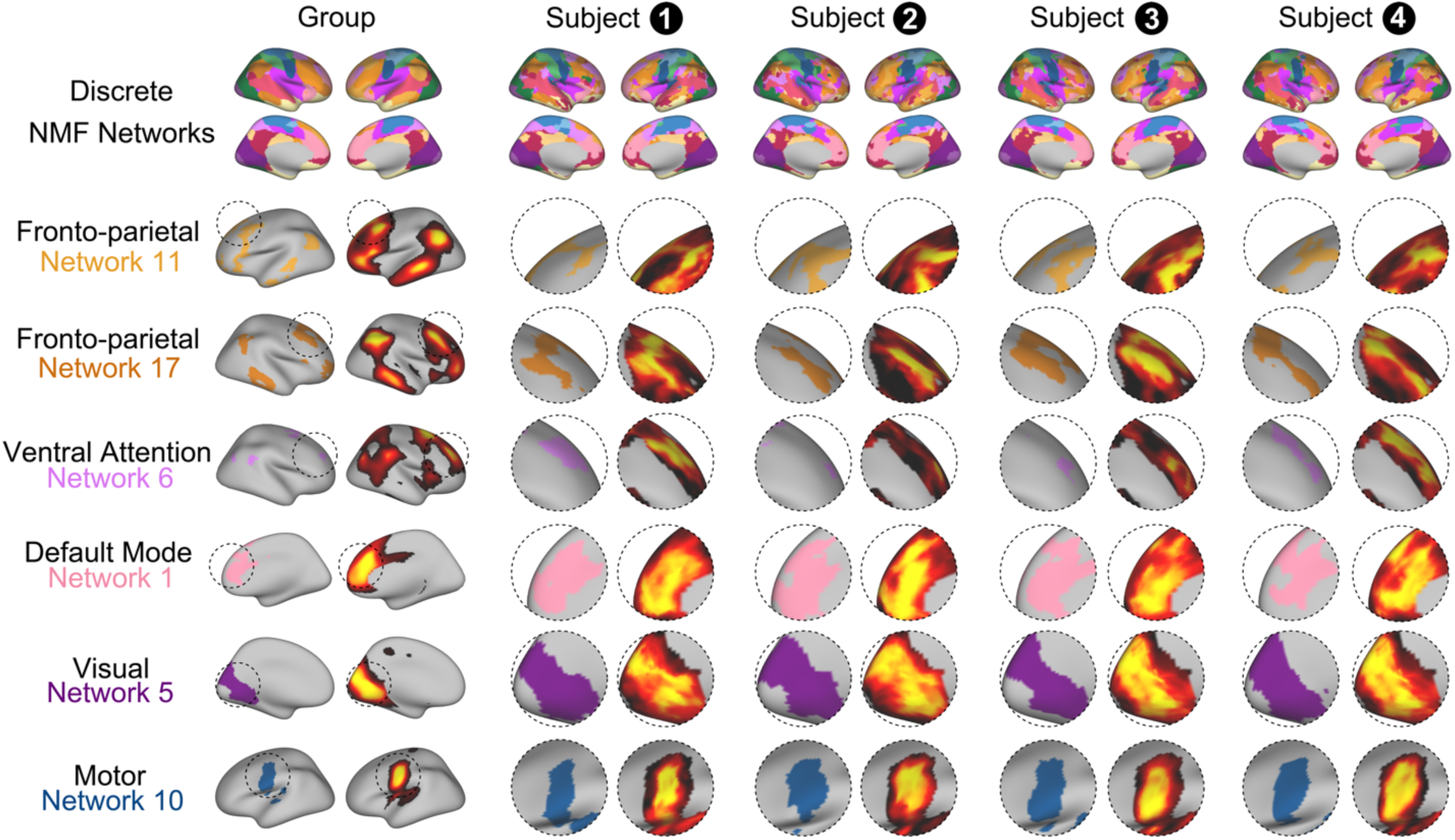
Individual subjects display distinct functional network topography. While the gross spatial distribution of networks was consistent across participants, distinct person-specific topographic features could be readily observed. Heterogeneity in the spatial distribution of networks was particularly apparent in higher-order networks including fronto-parietal, ventral attention, and default mode networks. In contrast, subject level representations of somatomotor and visual networks appeared to be much more consistent across individuals.

In order to evaluate this observation in the entire sample, we quantified the across-subject variance in network loadings using a non-parametric statistic (the median absolute deviation). As expected from prior reports in adults (Gordon et al., 2017b; Gordon et al., 2017c; Kong et al., 2018; Li et al., 2019; Mueller et al., 2013; Wang et al., 2015), we observed the highest across-subject variability in frontal, parietal, and temporal cortex (**Figure 4A**). When variability was ranked by network, we found that fronto-parietal networks had the highest topographic variability across subjects, whereas sensory and motor networks had the lowest (**Figure 4B**). Results were highly similar when the variability of discrete networks derived using NMF or MS-HBM were examined (**Supplementary Figure 3**). Having confirmed that functional topography is most variable in higher-order networks, we next evaluated whether this variation was related to brain maturation during youth.

**Figure 4.**
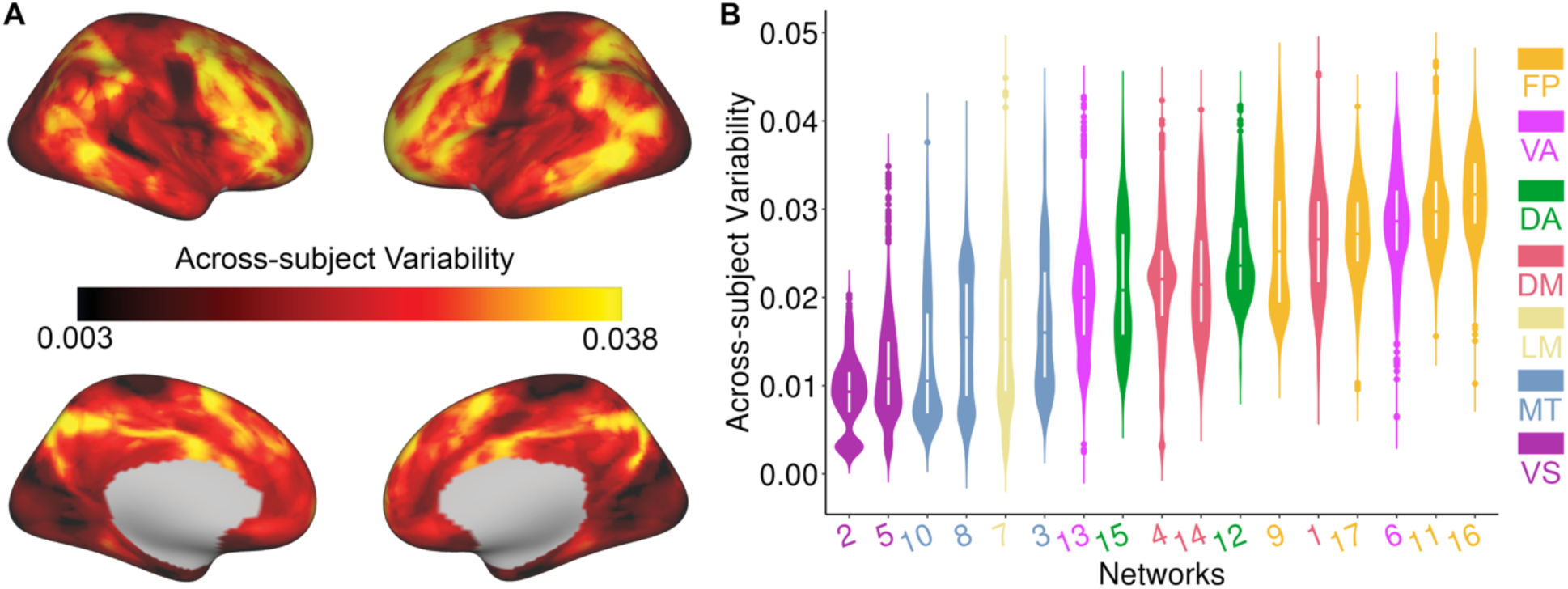
Variability of functional network topography is highest in executive networks. (**A**) A non-parametric measure of variability (median absolute deviation) revealed that functional topography was most variable across individuals in fronto-parietal cortex and least variable in visual and motor cortex. (**B**) Summarizing variability by network revealed that across-subject variability was highest in networks critical for executive functioning including fronto-parietal control networks and the ventral attention network. FP: fronto-parietal; VA: ventral attention; DA: dorsal attention; DM: default mode; LM: Limbic; MT: motor; VS: visual.

### Functional topography is refined with age and encodes brain maturity

As an initial step, we examined whether the total cortical representation of each network was associated with age. Specifically, for each network, we summed the loadings of all vertices to summarize the total cortical representation of each probabilistic network. Notably, as networks were derived in template space, this measure controls for individual differences in total surface area, which varies across development (Tamnes et al., 2017). As brain development is known to be a nonlinear process (Blakemore, 2012; Grayson and Fair, 2017; Tamnes et al., 2017), we used general additive models (GAMs; Wood (2004)) to capture both linear and nonlinear associations with age. Within each GAM, age was modeled using a penalized spline, while covarying for sex and in-scanner motion. After correcting for multiple comparisons with the Bonferroni method, these analyses revealed that the cortical representation of the limbic network (network 7, *Z* = 5.42, *P*_Bonf_ = 1.03 × 10^−6^, partial *r* = 0.20, Confidence Interval (CI) = [0.13, 0.27]) significantly increased with age, while that of the visual network (network 5, *Z* = −3.95, *P*_Bonf_ = 1.31 × 10^−3^, partial *r* = − 0.15, Confidence Interval (CI) = [−0.22, −0.08]) significantly decreased with age (**Figure 5A**).

**Figure 5.**
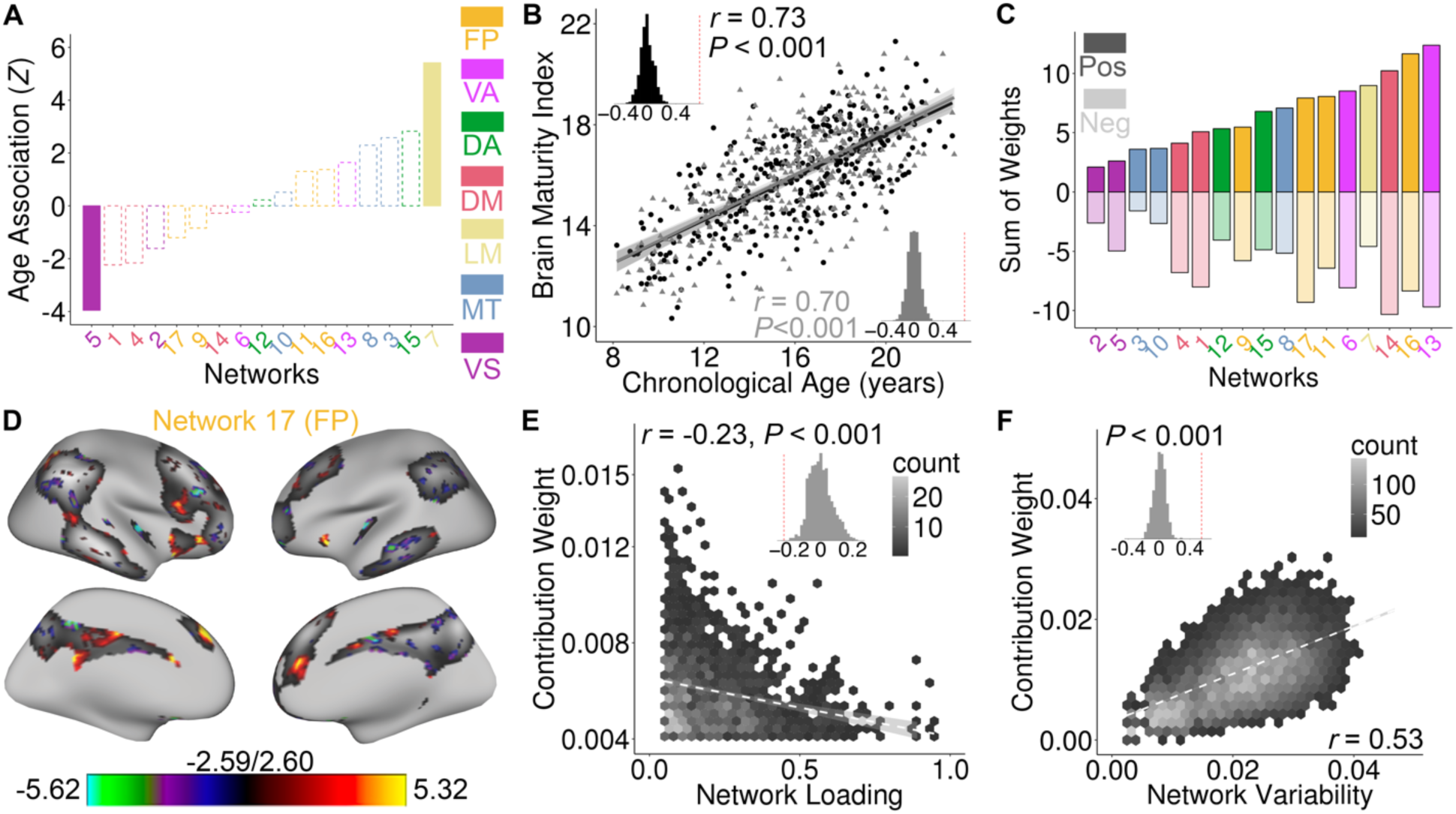
Functional topography evolves with age in youth and predicts unseen individuals’ brain maturity. (**A**) The total cortical representation of the limbic network increased with age, while the representation of the lateral visual network declined with age (*P*_Bonf_ < 0.05; dashed lines indicate networks with non-significant age effects). (**B**) Beyond this coarse summary measure, the complex pattern of developmental reconfiguration of functional topography could be used to predict age in unseen data using a multivariate ridge regression model with 2-fold cross-validation and nested parameter tuning. Data points represent predicted age of subjects in a model trained on independent data; inset histogram represents the null distribution of prediction accuracy from a permutation test. (**C**) Examining the sum of the absolute model weight of all vertices within each network revealed that high-order networks in association cortex contributed the most to predicting age. Both positive and negative associations with age were present within each network. (**D**) Model feature weights driving prediction were highest at network edges; the 25% vertices of network 17 that had the highest absolute contribution weight are displayed. (**E**) Absolute feature weight was negatively correlated with network loadings across vertices for network 17; inset displays spatial association compared to null distribution from spin test. (**F**) Functional network maturation is constrained by network variability. Vertices that contributed the most to the multivariate age prediction model were those that varied most across subjects. FP: fronto-parietal; VA: ventral attention; DA: dorsal attention; DM: default mode; LM: Limbic; MT: motor; VS: visual.

It should be noted that a coarse summary measure such as the total network representation does not capture complex patterns of topographic reconfiguration. However, mass-univariate models of each network across individual verticies are limited by necessary corrections for multiple comparisons, and cannot model multivariate relationships within high-dimensional data (see **Supplementary Figure 4**). Accordingly, we used a multivariate approach to understand the degree to which the overall pattern of functional topography encoded developmental information. Specifically, we used ridge regression with nested two-fold cross validation (2F-CV, **Supplementary Figure 5**) to predict the age of an unseen individual based on the functional topography of all networks. Using training and testing sets that were matched on age, we calculated both the mean absolute error (MAE) and the partial correlation between the predicted age (“brain age”) and chronologic age in the test set, while controlling for sex and motion. Model significance was evaluated using permutation testing, where the correspondence between training subjects’ network topography and their age was shuffled at random. This multivariate analysis revealed that the complex pattern of network topography could accurately predict an unseen individual’s age with a high degree of accuracy (**Figure 5B**): the partial correlation between the predicted age and chronological age was 0.73 (*P_perm_* < 0.001), while MAE was 1.84 years (*P_perm_* < 0.001). We repeated this procedure while reversing the training and testing sets, and found very similar results (partial *r* = 0.70, *P_perm_* < 0.001; MAE = 1.89 years, *P_perm_* < 0.001).

To understand the developmental effects underlying these results, we evaluated model feature weights. In the multivariate model, each vertex received a feature weight for each network. Summing the absolute weights within each network, we found that high-order association networks contributed the most to the multivariate model (**Figure 5C**). However, we also found that there was a complex pattern of both positive and negative relationships with age (**Supplementary Figure 6**), cohering with the initial finding that the total network representation did not change with age in most networks. Examining the spatial distribution of these feature weights, we observed that vertices with the highest weights often tended to be at the edge of the network. For example, refinement of network boundaries with age was particularly prominent in several fronto-parietal networks (e.g., network 17; **Figure 5D**). In order to quantify this observation, we examined the relationship between mean network loading and the absolute weight of features in the multivariate model predicting age. Consistent with a process of edge refinement and spatial differentiation between networks, we found that higher feature weights were present at edge vertices with low loadings in multiple networks, including fronto-parietal networks (**Figure 5E** and **Supplementary Figure 7**).

As a final step, we sought to understand whether variability of functional topography constrained patterns of network maturation. To concisely summarize the spatial contribution of locations in the multivariate model, we summed the absolute weights of each vertex across networks, and related this to our non-parametric measure of network variability (see **Figure 4A**). We found that multivariate patterns of brain maturation were driven by vertices with high across-participant variability, and were present primarily in frontal, parietal, and temporal cortex (**Figure 5F**; *r* = 0.53, *P_spin_* < 0.001).

We conducted several supplementary analyses to confirm that our results were robust to methodological choices. In order to ensure that our matched split of the data was representative, we repeated this procedure with 100 random splits of the data, which returned highly consistent results and feature weights (mean partial *r* = 0.69, *P_perm_* < 0.001; mean MAE = 1.93 years, *P_perm_* < 0.001; **Supplementary Figure 8**). For comparison, we repeated this procedure using both the discrete network parcellation derived from NMF and also that from MS-HBM. While still highly significant (*P_perm_* < 0.001), not considering network probability mildly degraded predictive accuracy (see **Supplementary Figure 8**). Taken together, these results demonstrate that functional network topography encodes brain maturity, is driven by refinement of higher-order association networks, and is constrained by the individual variability of these systems.

### Control network topography predicts individual differences in executive function

Having found that functional topography accurately encoded brain maturation, we next evaluated the implications of topographic variability for cognition. Specifically, we investigated whether variation in functional network topography predicted individual differences in executive function. Executive function was summarized using a previously-published factor analysis of the Penn Computerized Neurocognitive Battery (Moore et al., 2015). While controlling for age, sex, and motion, general additive models revealed that the improved executive performance was associated with a greater total cortical representation of bilateral fronto-parietal control networks and the cingulo-opercular ventral attention network (**Figure 6A;** left fronto-parietal: network 11, *Z* = 5.88, *P*_Bonf_ = 1.09 × 10^−7^, partial *r* = 0.22, CI = [0.15, 0.29]; right fronto-parietal: network 17, *Z* = 5.23, *P*_Bonf_ = 3.90 × 10^−6^, partial *r* = 0.19, CI = [0.12, 0.26]; ventral attention: network 6, *Z* = 4.41, *P*_Bonf_ = 2.06 × 10^−4^, partial *r* = 0.17, CI = [0.09, 0.24]). In contrast, greater representation of the right temporal default mode network (network 4, *Z* = −3.43, *P*_Bonf_ = 0.01, partial *r* = −0.13, CI = [−0.20, −0.06]) and the limbic network (network 7, *Z* = −3.35, *P*_Bonf_ = 0.01, partial *r* = −0.13, CI = [−0.20, −0.05]) were associated with reduced executive performance. High resolution analyses of vertices provided convergent results (**Supplementary Figure 9**).

**Figure 6.**
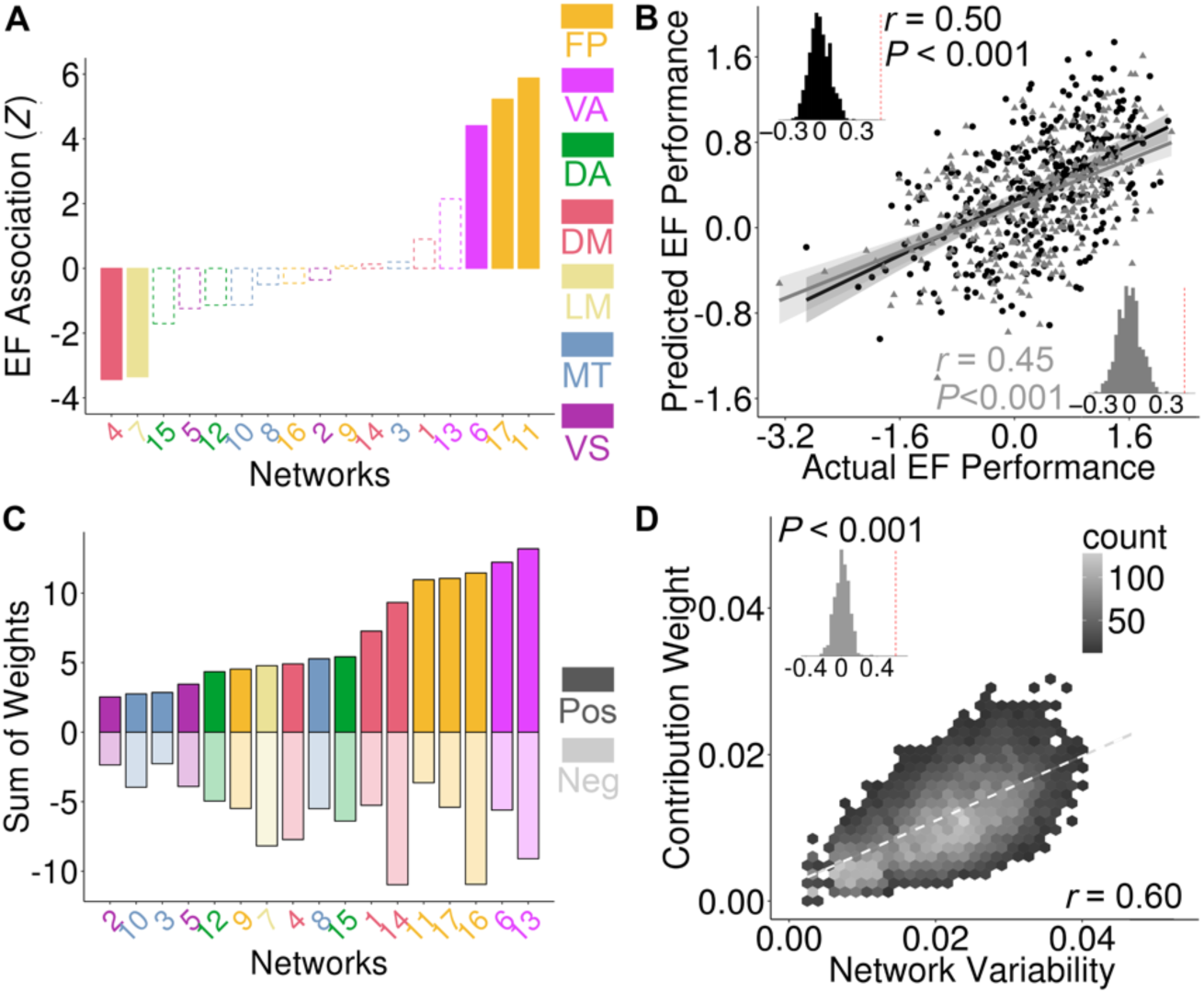
Functional topography of control networks predicts individual differences in executive function. (**A**) Executive performance was positively correlated with the total cortical representation of two fronto-parietal networks and one ventral attention network (*P*_Bonf_ < 0.05; dashed lines indicates networks with non-significant age effects). (**B**) The complex pattern of functional network topography predicted executive function in unseen data using a multivariate ridge regression model with 2-fold cross-validation and nested parameter tuning (data points represent predicted age of subjects by a model trained on independent data; inset histogram represents the distribution of prediction accuracy from a permutation test). (**C**) The most important topographic features in this model were found in association cortex critical for executive functioning, and were maximal in the fronto-parietal control network and the ventral attention network. (**D**) The vertices that contributed the most in this multivariate model were those that varied most across participants. EF: executive function; FP: fronto-parietal; VA: ventral attention; DA: dorsal attention; DM: default mode; LM: Limbic; MT: motor; VS: visual.

As for our analyses of development, we also evaluated the degree to which an individual’s multivariate pattern of network topography could be used to predict executive performance using a model trained on independent data. We found that an individualized functional topography accurately predicted executive functioning in matched split-half samples while controlling for age, sex and motion (**Figure 6B**; split 1: partial *r* = 0.50, MAE = 0.57, *P_perm_* < 0.001; split 2: partial *r* = 0.45, MAE = 0.59, *P_perm_* < 0.001). Critically, topographic features within the ventral attention and fronto-parietal networks were the most predictive of individual differences in executive functioning (**Figure 6C & Supplementary Figure 10**). Multivariate feature weights aligned with analyses of the total network representation, with a preponderance of positive relationships with executive performance being found within executive networks. As for patterns of brain maturation, we found that the topographic features that predicted executive functioning were those that varied most across individuals (**Fig. 7C**; *r* = 0.60, *P_spin_* < 0.001).

**Figure 7.**
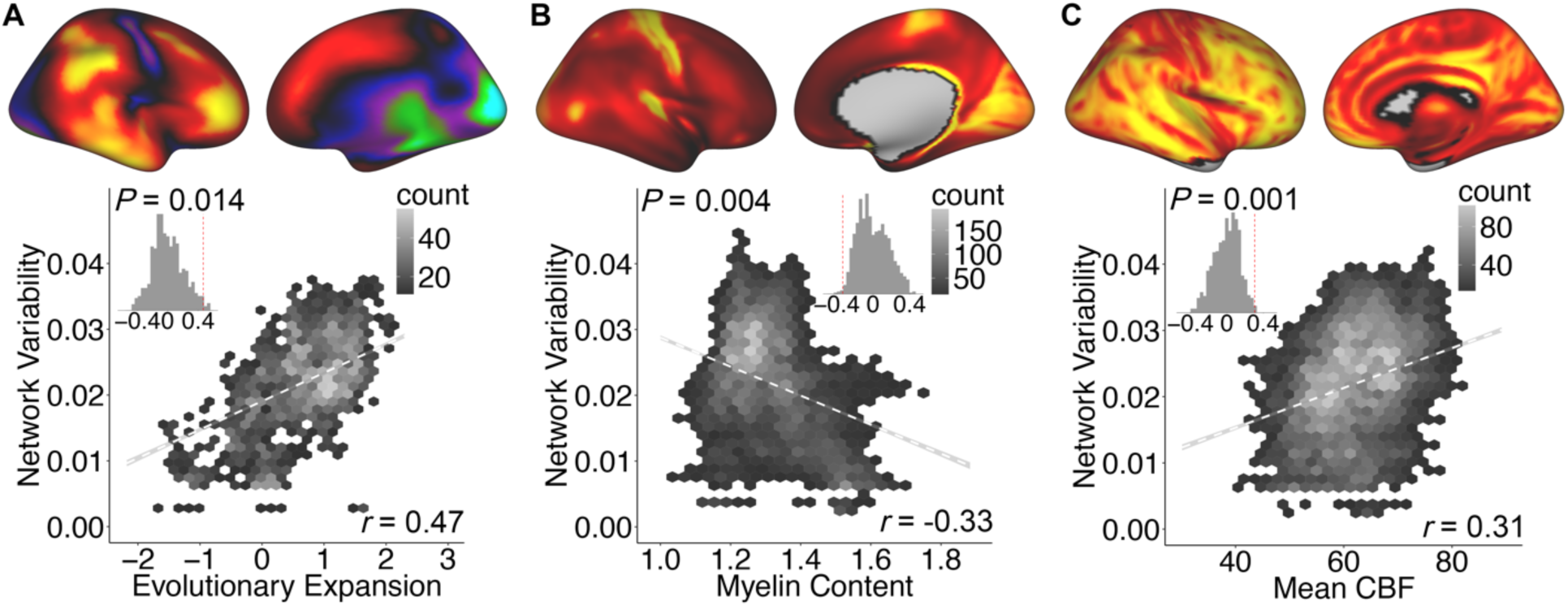
Variability in functional topography aligns with fundamental properties of brain organization. Higher variability in network topography was associated with greater evolutionary expansion (**A**), lower myelin content (**B**), and higher cerebral blood flow (**C**). Inset histograms represent spatial association compared to a null distribution obtained from spatial permutation testing.

To again ensure our initial matched split of the data was representative, we repeated the analysis with 100 random splits, which yielded highly consistent results (mean partial *r* = 0.47, mean MAE = 0.58; **Supplementary Figure 11**). Furthermore, we repeated this procedure using a discrete network parcellation from either NMF or MS-HBM, which returned similar results (**Supplementary Figure 11**). Finally, we tested the specificity of associations with executive function. We found that the ability of functional topography to predict either memory (split 1: partial *r* = 0.23, *P_perm_* = 0.002, MAE = 0.70, *P_perm_* < 0.001; split 2: partial *r* = 0.23, *P_perm_* = 0.008, MAE = 0.69, *P_perm_* < 0.001) and social cognition (split 1: partial *r* = 0.11, *P_perm_* = 0.07, MAE = 0.63, *P_perm_* < 0.001; split 2: partial *r* = 0.09, *P_perm_* = 0.16, MAE = 0.62, *P_perm_* < 0.001) was substantially lower than the executive function. These results emphasize that patterns of functional topography within high-order control networks that are most variable across individuals predict individual differences in executive function.

### Variability of functional topography is constrained by fundamental properties of brain organization

Having demonstrated that variability in functional topography predicts both brain maturation and individual differences in executive function, we next sought to understand if topographic variability is constrained by evolutionary properties of brain structure. One prominent theory of cortical organization suggests that large-scale association networks arose in evolution by becoming untethered from rigid developmental programming present in lower-order somatosensory systems (Buckner and Krienen, 2013). This theory is supported by the distribution of cortical myelin: association cortex that has undergone dramatic evolutionary expansion also has greatly reduced myelination compared to somatosensory cortex. Notably, such lightly-myelinated association cortex has higher metabolic demands and receives a disproportionate proportion of cerebral blood flow. Having demonstrated that functional topography is the highest in association cortex, and that this variation predicts both age and executive function, we sought to directly relate topographic variability to these fundamental properties of brain organization. Specifically, we hypothesized that higher variability in functional topography would be associated with greater evolutionary expansion, reduced myelin content, and higher cerebral blood flow.

Using our statistically conservative spatial permutation testing procedure, we found that the cortical regions that exhibit the most topographic variability are also those that have expanded the most in evolution (**Figure 8A**; *r* = 0.47, *P_spin_* = 0.014). In contrast, variability in cortical topography was inversely related to cortical myelin content (**Figure 8B**; *r* = −0.33, *P_spin_* = 0.004). Finally, higher network variability was significantly associated with cerebral blood flow (**Figure 8C**, *r* = 0.31, *P_spin_* = 0.001). Thus, lightly-myelinated cortex that has expanded dramatically in evolution and receives a disproportion degree of cerebral blood flow also exhibits the greatest variability in functional neuroanatomy during youth.

## DISCUSSION

In this study we leveraged advances in machine learning and a large sample of youth to delineate how the functional topography of the cortex develops during youth and supports executive function. Building upon findings from studies of adults, we confirmed that networks necessary for executive function also show the most topographic variability in childhood and adolescence. Critically, we demonstrate that these same networks are refined during development and predict individual differences in executive performance. Finally, we provide evidence that topographic variability is strongly linked to fundamental properties of brain organization. Taken together, these results offer a new account of both developmental plasticity and diversity, and highlight potential for progress in personalized diagnostics and therapeutics.

This work builds on a series of studies that have documented inter-individual variability in the spatial layout of canonical functional networks (Braga and Buckner, 2017; Glasser et al., 2016; Gordon et al., 2017a; Gordon et al., 2017c; Kong et al., 2018; Laumann et al., 2015; Li et al., 2017; Li et al., 2019; Wang et al., 2015; Wang et al., 2018). Though previous efforts have used a variety of analysis techniques to define functional networks in individuals, they have yielded convergent results. Prior work in adults has emphasized that variability in functional topography is heterogeneously distributed across the cortex, with higher-order functional networks responsible for control processes displaying the greatest variance across individuals (Braga and Buckner, 2017; Gordon et al., 2017b; Gordon et al., 2017c; Kong et al., 2018; Li et al., 2019; Wang et al., 2015). Building upon these results from adults, we found that these same higher-order networks are also the most variable in youth. Moreover, we demonstrated that this variability in functional topography constrains patterns of brain maturation and is associated with individual differences in executive capability during youth.

Our results demonstrate that individual variation in functional network topography is linked to both brain development and executive functioning. Specifically, we found that at any given age a greater cortical representation of control networks is associated with improved executive performance. In contrast, while the relative proportion of cortex allocated to association networks does not appear to undergo large shifts in youth, machine learning techniques were able to decode developmental processes from the complex pattern of functional topography. Using multivariate ridge regression, functional topography predicted an unseen individual’s age with a high degree of accuracy, with association networks contributing the most to this prediction.

When integrated across levels of analysis, these developmental results are consistent with a process of network differentiation (Baum et al., 2017; Fair et al., 2007a; Gu et al., 2015; Sherman et al., 2014; Wig, 2017). For example, while the fronto-parietal, ventral attention, and default mode networks did not change in its total cortical representation with age, examination of multivariate feature weights revealed a complex pattern of reconfiguration with both positive and negative associations with age. Maturational changes were frequently concentrated at network boundaries, suggesting that the network borders are refined in development. These results are align with the protracted process of network differentiation within higher-order cortex, whereby functional systems with divergent cognitive roles (such as executive and the default mode networks) become more distinct in their functional topography (Baum et al., 2017; Fair et al., 2007a; Sherman et al., 2014). This process may partially explain previous reports of developmental network segregation, which is among the most-replicated results in developmental cognitive neuroscience (Baum et al., 2017; Fair et al., 2007a; Gu et al., 2015; Satterthwaite et al., 2012; Sherman et al., 2014; Wig, 2017). Increasingly differentiated functional networks would reduce mixing of signals between networks, and thus appear to have reduced inter-network connectivity. Indeed, this possibility was recently suggested by work in adults (Bijsterbosch et al., 2018; Li et al., 2019), which showed that topography and connectivity have distinct contributions to individual differences, and that ignoring topography aliases topographic signals into measurement of connectivity. Although further research is needed in parallel human and animal models, the observed developmental network refinement may be in part driven by focused myelination and ongoing pruning, which continues in association cortex through early adulthood (Hagmann et al., 2010).

In contrast to such complex network differentiation, we found that youth with a greater cortical representation of executive control networks had better executive functioning. Results were convergent at all scales of analysis, including network summary measures, high-resolution analyses, and integrative multivariate models. These data suggest that the marked between-subject variability of executive network topography has implications for behavior, and may be relevant for neuropsychiatric disorders. At present, the origins of these individual differences in executive network topography remain unknown. However, the substantial heritability of both cognitive performance and functional connectivity suggests that topography is at least in part genetically encoded (Colclough et al., 2017; Mollink et al., 2019). Furthermore, accruing evidence from animal models and translational studies in humans emphasizes the likely importance of in-utero and early-life stressors, which could potentially impact developmental initialization of functional network topography (Graham et al., 2019). In the future, it will be possible to test this hypothesis using a combination of studies in animal models and human infants with varying levels of stress exposure.

The topographic variability, developmental plasticity, and potential vulnerability of higher-order control networks may be in part understood by evolutionary constraints. Leveraging independent data from multiple sources, we found that variability and developmental change in topography is maximal in the same executive networks that have undergone the most evolutionary expansion (Hill et al., 2010). These networks have low myelin content (Glasser and Van Essen, 2011) and receive the greatest relative blood flow (Satterthwaite et al., 2014c). As noted in multiple prior accounts of cortical organization, higher order executive networks are spatially embedded between somatosensory and default mode regions (Margulies et al., 2016). One prominent theory suggests that these systems may have become untethered from the detailed developmental programing of highly conserved somatosensory cortex as part of their rapid evolutionary expansion, thus allowing for non-canonical circuit properties and enhanced individual variability (Buckner and Krienen, 2013). Our results are consistent with such an account, as highly variable cortical networks that are under diminished anatomic constraints also evince the most marked developmental change and individual variability.

Several potential limitations and countervailing strengths of the present study should be noted. First, all data presented here were cross-sectional, which precludes inference regarding within-individual developmental effects. Ongoing follow-up studies will yield informative longitudinal data, as will large-scale studies such as the Adolescent Brain and Cognitive Development study (Casey et al., 2018). Second, we used data combined across three fMRI runs, including two where an fMRI task was regressed from the data. This choice was motivated by convergent results from several independent studies, which have shown that functional networks are primarily defined by individual-specific rather than task-specific factors (Gratton et al., 2018) and that intrinsic networks during task performance are similar to those present at rest (Fair et al., 2007b). By including task-regressed data, we were able to generate individualized networks using 27-minutes of high quality data. Prior work suggests that parcellations created using a timeseries of this length show high concordance (r~0.92) with those generated using 380 minutes of data (Laumann et al., 2015), and that longer timeseries allow for greatly improved prediction of individual differences (Elliott et al., 2019). Third, it should be acknowledged that our individualized parcellations are data driven, and at present there are no techniques for ascertaining neurobiological ground truth in humans. Nonetheless, it is reassuring that our results were robust to substantial methodological variation, including the use of a completely independent method for defining individualized network. Fourth, because children tend to move more during the scanning session, in-scanner motion continues to be a concern for all functional imaging studies of brain development. However, in this study we rigorously followed best practices for mitigating this confound, including use of an extensively-benchmarked, top-performing preprocessing pipeline and co-varying for motion in all hypothesis testing (Ciric et al., 2018; Satterthwaite et al., 2013a). Use of these conservative procedures bolsters confidence that our observed results are not driven by the confounding influence of in-scanner motion.

These limitations notwithstanding, we provide novel evidence that individual-specific functional network topography is refined during development and supports executive function. These findings also emphasize the relevance of functional network topography for translational clinical neuroscience. Notably, higher order networks that undergo the most developmental change also are the same networks that have been linked to diverse neuropsychiatric illnesses including psychosis, mood disorders, and ADHD (Cole et al., 2014; Xia et al., 2018). As neuropsychiatric conditions are increasingly conceptualized as disorders of brain development (Insel, 2014), functional topography may be critically important for understanding the neurodevelopmental substrates of these debilitating disorders, and allow for early identification and intervention in youth at risk. Finally, these results suggest clear next steps for integration with clinical trials of personalized neuromodulatory interventions that are targeted using the specific functional neuroanatomy of an individual patient.

## ACKNOWLEDGEMENTS

Thanks to Ru Kong and Jingwei Li for guidance in implementating the MS-HBM method. This study was supported by grants from National Institute of Mental Health: R01MH113550 (T.D.S. & D.S.B.), R01EB022573 (C.D. & Y.F.), R01MH107703 (T.D.S.), RF1MH116920 (D.J.O., T.D.S. & D.S.B.), and R01MH112847 (R.T.S. & T.D.S.). The PNC was supported by MH089983 and MH089924. Additional support was provided by the Lifespan Brain Institute at Penn and the Children’s Hospital of Philadelphia, by P50MH096891 to R.E.G., R01MH11186 to D.J.O., K01MH102609 to D.R.R., R01MH107235 to R.C.G., R01NS085211 to R.T.S., and the Dowshen Program for Neuroscience. The content is solely the responsibility of the authors and does not represent the official views of any of the funding agencies. Thanks to Chad Jackson for data and systems support, *in memoriam*.

## AUTHOR CONTRIBUTIONS

T.D.S and Z.C. designed the research; Z.C. and T.D.S. performed the analyses with support from H.L., B.L., A.A., G.L.B., M.C.; H.L. and Y.F. provided parcellation tools; H.L. replicated the analysis; Z.C., C.H.X., T.D.S made figures; A.A., M.C., T.M.M, D.R.R, T.D.S. completed data preprocessing; R.E.G, R.C.G, C.D and T.D.S. provided resources; D.S.B., A.A-B, D.J.O., R.T.S., D.H.W., C.D. D.A.F, Y.F. provided critical comments on analysis; Z.C. and T.D.S wrote the manuscript, with contributions from B.L. and comments from all other authors.

## METHOD DETAILS

### Participants

Overall, 1,601 participants were studied as part of the PNC (Satterthwaite et al., 2014a). However, 340 subjects were excluded due to clinical factors, including medical disorders that could affect brain function, current use of psychoactive medications, prior inpatient psychiatric treatment, or an incidentally encountered structural brain abnormality. Among the 1,261 subjects eligible for inclusion, 54 subjects were excluded for a low quality T1-weighted image or low quality FreeSurfer reconstructions. Of the 1,207 subjects with a usable T1 image and adequate FreeSurfer reconstruction, 514 participants were excluded for missing functional data or poor functional image quality; all participants were required to have three functional runs which passed QA. Specifically, as in prior work (Ciric et al., 2018), a functional run was excluded if mean relative RMS framewise displacement was higher than 0.2mm, or it had more than 20 frames with motion exceeding 0.25mm. This set of exclusion criteria resulted in a final sample of 693 participants (**Supplementary Figure 1**), with mean age 15.93 years, SD = 3.23 years; the sample included 301 males and 392 females. All study procedures were approved by the Institutional Review Boards of both the University of Pennsylvania and the Children’s Hospital of Philadelphia.

### Cognitive assessment

The Penn computerized neurocognitive battery (Penn CNB) was administered to all participants as part of a session separate from neuroimaging. The CNB consists of 14 tests adapted from tasks applied in functional neuroimaging to evaluate a broad range of cognitive domains (Gur et al., 2012). These domains include executive function (abstraction and mental flexibility, attention, working memory), episodic memory (verbal, facial, spatial), complex cognition (verbal reasoning, nonverbal reasoning, spatial processing), social cognition (emotion identification, emotion differentiation, age differentiation), and sensorimotor and motor speed. Accuracy for each test was *z*-transformed. As previously described in detail, factor analysis was used to summarize these accuracy scores into three factors (Moore et al., 2015), including executive and complex cognition, episodic memory, and social cognition. Here, we focused on the executive and complex cognition factor score. However, episodic memory and social cognition factor scores were evaluated in specificity analyses.

### Image acquisition

As previously described (Satterthwaite et al., 2014a), all MRI scans were acquired using the same 3T Siemens Tim Trio whole-body scanner and 32-channel head coil at the Hospital of the University of Pennsylvania.

#### Structural MRI

Prior to functional MRI acquisitions, a 5-min magnetization-prepared, rapid acquisition gradient-echo T1-weighted (MPRAGE) image (TR = 1810 ms; TE = 3.51 ms; TI = 1100 ms, FOV = 180 × 240 mm^2^, matrix = 192 × 256, effective voxel resolution = 0.9 × 0.9 × 1 mm^3^) was acquired.

#### Functional MRI

We used one resting-state and two task-based (i.e., *n*-back and emotion identification) fMRI scans as part of this study. All fMRI scans were acquired with the same single-shot, interleaved multi-slice, gradient-echo, echo planar imaging (GE-EPI) sequence sensitive to BOLD contrast with the following parameters: TR = 3000 ms; TE = 32 ms; flip angle = 90°; FOV = 192 × 192 mm^2^; matrix = 64 × 64; 46 slices; slice thickness/gap = 3/0 mm, effective voxel resolution = 3.0 × 3.0 × 3.0 mm^3^. Resting-state scans had 124 volumes, while the *n*-back and emotion recognition scans had 231 and 210 volumes, respectively. Further details regarding the *n*-back (Satterthwaite et al., 2013b) and emotion recognition (Wolf et al., 2015) tasks have been described in prior publications.

#### Field map

In addition, a B0 field map was derived for application of distortion correction procedures, using a double-echo, gradient-recalled echo (GRE) sequence: TR = 1000ms; TE1 = 2.69ms; TE2 = 5.27ms; 44 slices; slice thickness/gap = 4/0 mm; FOV = 240 mm; effective voxel resolution = 3.8×3.8×4 mm.

#### Scanning procedure

Before scanning, to acclimate subjects to the MRI environment, a mock scanning session where subjects practiced the task was conducted using a decommissioned MRI scanner and head coil. Mock scanning was accompanied by acoustic recordings of the noise produced by gradient coils for each scanning pulse sequence. During these sessions, feedback regarding head movement was provided using the MoTrack motion tracking system (Psychology Software Tools). Motion feedback was given only during the mock scanning session. To further minimize motion, before data acquisition, subjects’ heads were stabilized in the head coil using one foam pad over each ear and a third over the top of the head.

### Image processing

The structural images were processed using FreeSurfer (version 5.3) to allow for the projection of functional timeseries to the cortical surface (Fischl, 2012). Functional images were processed using a top-performing preprocessing pipeline implemented using the eXtensible Connectivity Pipeline (XCP) Engine (Ciric et al., 2018). This pipeline included (1) correction for distortions induced by magnetic field inhomogeneity using FSL’s FUGUE utility, (2) removal of the initial volumes of each acquisition (i.e., 4 volumes for resting-state fMRI and 6 volumes for emotion recognition task fMRI), (3) realignment of all volumes to a selected reference volume using FSL’s MCFLIRT, (4) interpolation of intensity outliers in each voxel’s time series using AFNI’s 3dDespike utility, (5) demeaning and removal of any linear or quadratic trends, and (6) co-registration of functional data to the high-resolution structural image using boundary-based registration. Images were de-noised using a 36-parameter confound regression model that has been shown to minimize associations with motion artifact while retaining signals of interest in distinct sub-networks. This model included the six framewise estimates of motion, the mean signal extracted from eroded white matter and cerebrospinal fluid compartments, the mean signal extracted from the entire brain, the derivatives of each of these nine parameters, and quadratic terms of each of the nine parameters and their derivatives. Both the BOLD-weighted time series and the artefactual model time series were temporally filtered using a fist-order Butterworth filter with a passband between 0.01 and 0.08 HZ to avoid mismatch in the temporal domain (Hallquist et al., 2013). Furthermore, to derive “pseudo-resting state” timeseries that were comparable across runs, the task activation model was regressed from n-back or emotion identification fMRI data (Fair et al., 2007b). The task activation model and nuisance matrix were regressed out using AFNI’s 3dTproject.

For each modality, the fMRI timeseries of each individual were projected to each subject’s FreeSurfer surface reconstruction and smoothed on the surface with a 6-mm full-width half-maximum (FWHM) kernel. The smoothed data was projected to the *fsaverage5* template, which has 10,242 vertices on each hemisphere (18,715 vertices in total after removing the medial wall). Finally, we concatenated the three fMRI acquisitions, yielding timeseries of 27 minutes, 45 seconds (555 timepoints) in total.

### Regularized non-negative matrix factorization

As previously described in detail (Li et al., 2017), we used non-negative matrix factorization (NMF) (Lee and Seung, 1999) to derive individualized functional networks. The NMF method factors the data by positively weighting cortical elements that covary, leading to a highly specific and reproducible parts-based representation (Lee and Seung, 1999; Sotiras et al., 2017). Our approach was enhanced by a group consensus regularization term that preserves the inter-individual correspondence, as well as a data locality regularization term that makes the decomposition robust to imaging noise, improves spatial smoothness, and enhances functional coherence of the subject-specific functional networks (see Li et al. (2017) for details of the method; see also: https://github.com/hmlicas/Collaborative_Brain_Decomposition). As NMF requires the input to be nonnegative values, we re-scaled the data by shifting time courses of each vertex linearly to ensure all values were positive (Li et al., 2017). To avoid features in greater numeric ranges dominating those in smaller numeric range, we further normalized the time course by its maximum value so that all the time points have values in the range of [0, 1].

Given a group of *n* subjects, each having fMRI data *X^i^* ∈ *R*^×^, *i* = 1, …, *n*, consisting of *S* vertices and *T* time points, we aimed to find *K* non-negative functional networks 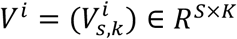 and their corresponding time courses 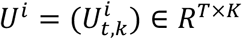 for each subject, such that

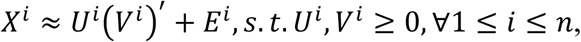

where (*V^i^*)′ is the transpose of (*V^i^*), and *E^i^* is independently and identically distributed (i.i.d) residual noise following Gaussian distribution with a probability density function of 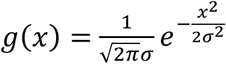. Both *U^i^* and *V^i^* were constrained to be non-negative so that each functional network does not contain any anti-correlated functional units (Lee and Seung, 1999). A group consensus regularization term was applied to ensure inter-individual correspondence, which was implemented as a scale-invariant group sparsity term on each column of *V^i^* and formulated as

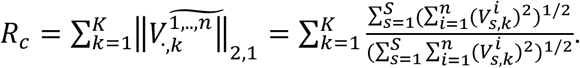

The data locality regularization term was applied to encourage spatial smoothness and coherence of the functional networks using graph regularization techniques (Cai et al., 2011). The data locality regularization term was formulated as

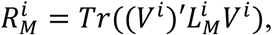

where 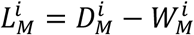 is a Laplacian matrix for subject *I*, 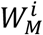 is a pairwise affinity matrix to measure spatial closeness or functional similarity between different vertices, and 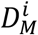 is its corresponding degree matrix. The affinity between each pair of spatially connected vertices (i.e., vertices *a* and *b*) was calculated as 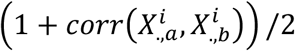, where 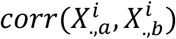 is the Pearson correlation coefficient between time series 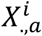 and 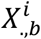, and others were set to zero so that 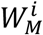, has a sparse structure. We identified subject specific functional networks by optimizing a joint model with integrated data fitting and regularization terms formulated by

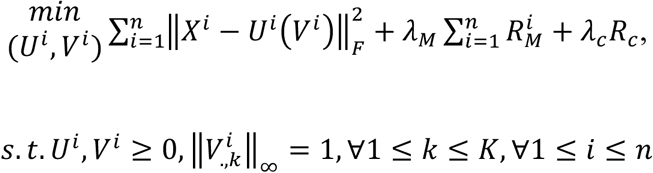

where 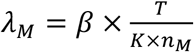 and 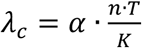 are used to balance the data fitting, data locality, and group consensus regularization terms, *n_M_* is the number of neighboring vertices, *α* and *β* are free parameters. For this study, we used identical paramters settings as in prior validation studies (Li et al., 2017).

### Defining individualized networks

Our approach for defining individualized networks included three steps (see **Figure 1**). In the first two steps, a consensus group atlas was created. In the third step, this group atlas was used to define individualized networks for each participant. We decomposed the whole-brain into 17 networks, which allowed for a direct comparison to other methods used in prior work (Kong et al., 2018; Wang et al., 2015; Yeo et al., 2011).

#### Step 1: Group network initialization

Although individuals exhibit distinct network topography, they are also broadly consistent (Gordon et al., 2017c; Gratton et al., 2018). Therefore, we first generated a group atlas and used it as an initialization for individualized network definition. In this way, we also ensured spatial correspondence across all subjects. This strategy has also been applied in other methods for individualized network definition (Kong et al., 2018; Wang et al., 2015). To avoid the group atlas being driven by outliers and to reduce the computational cost, a bootstrap strategy was utilized to perform the group-level decomposition multiple times on a subset of randomly selected subjects. Subsequently, the resulting decomposition results were fused to obtain one robust initialization that is highly reproducible. As previously (Li et al., 2017), we randomly selected 100 subjects and temporally concatenated their timeseries, resulting in a timeseries matrix with 55,500 rows (time-points) and 18,715 columns (vertices). Notably, the choice of sub-sample size did not impact results (sub-samples of 200 and 300 were also evaluated). We applied the above-mentioned regularized non-negative matrix factorization method with a random non-negative initialization to decompose this matrix (Lee and Seung, 1999). A group-level network loading matrix *V* was acquired, which had 17 rows and 18,715 columns. Each row of this matrix represents a functional network, while each column represents the loadings of a given cortical vertex. As previously (Li et al., 2017), this procedure was repeated 50 times, each time with a different subset of subjects; this yielded 50 different group atlases.

#### Step 2: Group network consensus

Next, we combined the 50 group network atlases to obtain one robust and highly reproducible group network atlas using spectral clustering (Li et al., 2017). Specifically, we concatenated the 50 group parcellations together across networks and acquired a matrix with 850 rows (i.e., functional networks, abbreviated as FN) and 18,715 columns (i.e., vertices). Inter-network similarity was calculated as

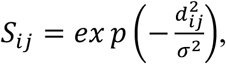

where *d_ij_* = 1 − *corr*(*FN_i_*, *FN_j_*), *corr*(*FN_i_*, *FN_j_*) is Pearson correlation coefficient between *FN_i_* and *FN_j_*, and *σ* is the median of *d_ij_* across all possible pairs of FNs. Then, we applied normalized-cuts (Cai et al., 2011) based spectral clustering method to group the 850 FNs into 17 clusters. For each cluster, the FN with the highest overall similarity with all other FNs within the same cluster was selected as the most representative. The final group network atlas was composed of the representatives of these 17 clusters.

#### Step 3: Individualized networks

In this final step, we derived each individual’s specific network atlas using NMF based on the acquired group networks (17 × 18,715 loading matrix) as initialization and each individual’s specific fMRI times series (555 × 18,715 matrix). See Li et al. (2017) for optimization details. This procedure yielded a loading matrix V (17 × 18,715 matrix) for each participant, where each row is a FN, each column is a vertex, and the value quantifies the extent each vertex belongs to each network. This probabilistic (soft) definition can be converted into discrete (hard) network definitions for display and comparison with other methods (Kong et al., 2018; Wang et al., 2015; Yeo et al., 2011) by labeling each vertex according to its highest loading.

### Multi-session Hierarchical Bayesian Model (MS-HBM)

To evaluate whether our results were robust to methodological variation, we also applied a a recently introduced multi-session hierarchical Bayesian model (MS-HBM, https://github.com/ThomasYeoLab/CBIG/tree/master/stable_projects/brain_parcellation/Kong2019_MSHBM) that has been used for defining individualized networks. See Kong et al. (2018) for the details of the method. Using a group atlas, this method calculates inter-subject resting-state functional connectivity (RSFC) variability, intra-subject RSFC variability, and finally parcellates for each single subject based on this prior information. We used the initialization values calculated using data from the Genomic Superstruct Project (GSP) dataset (Holmes et al., 2015), which were released along with Kong et al. (2018) (https://github.com/ThomasYeoLab/CBIG/tree/master/stable_projects/brain_parcellation/Kong2019_MSHBM/examples/input), as prior input of the single parcellation. Notably, the GSP was acquired using the identical fMRI sequences and scanning platform as the PNC. MS-HBM requires functional connectivity profiles of multiple sessions as input; here, the three fMRI runs were entered as three separate sessions. As in Kong et al. (Kong et al., 2018), we used MS-HBM to define 17 discrete individualized networks for each participant. Finally, we used the adjusted rand index (ARI) to calculate the similarity between the networks from MS-HBM and discretized networks from NMF.

### Spatial permutation testing (spin test)

In order to evaluate the significance of the alignment between individualized networks derived using NMF and MS-HBM, we used a spatial permutation procedure called the spin test (Alexander-Bloch et al., 2018; Gordon et al., 2016; Sotiras et al., 2017; Vandekar et al., 2015) (https://github.com/spin-test/spin-test). The spin test is a spatial permutation method based on angular permutations of spherical projections at the cortical surface. Critically, the spin test preserves the spatial covariance structure of the data and as such is far more conservative than randomly shuffling locations, which destroys the spatial covariance structure of the data and produce an unrealistically weak null distribution. In contrast, the spin test generates a null distribution of randomly rotated brain maps that preserve spatial features of the original map.

To evaluate the significance of the alignment between NMF and MS-HBM based networks, we compared the ARI of two parcellations to the ARI of 1,000 random rotations, generating a null distribution that preserves the spatial covariance structure. The permutation-based *p*-value was calculated as the proportion of times that the observed ARI was higher than the null distribution of ARI values from rotated parcellations. As described below, we also used the spin test to evaluate the significance of the alignment between across-subject parcellation variability to informative maps of brain organization.

### Homogeneity of functional networks

Network homogeneity is a commonly used method for evaluating the success of a functional parcellation (Gordon et al., 2016; Kong et al., 2018). As previously, network homogeneity was calculated as the average of the Pearson’s correlations between the time series of all pairs of vertices within each network (Kong et al., 2018). To summarize network homogeneity for comparisons across methods, we averaged the homogeneity value across networks.

### Across-subject variability of network topography

Prior studies of adults have consistently reported that across-subject variability of functional networks is high in higher-order association networks and lower in primary somatomotor and visual networks (Gordon et al., 2017b; Gordon et al., 2017c; Kong et al., 2018; Li et al., 2019; Mueller et al., 2013; Wang et al., 2015). Here we evaluated this observation in our sample of youth. For each of 17 networks, we calculated the median absolute deviation of loading values across all subjects for each vertex. We used this non-parametric measure of variance as loadings did not follow normal distribution. Next, we averaged the 17 median absolute deviation maps to generate the final across-network variability map that quantified the across-subject parcellation variance at each vertex.

Additionally, we also calculated the network variability of the discretized network atlas, allowing for further validation of our main results and better comparison to other methods (Kong et al., 2018). Specifically, we used entropy to define variability (Hoskisson et al., 1993):

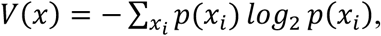

where *x* is a vertex; *x_i_* is a value of the vertex *x*, which has 17 values; *p(x_i_)* is the proportion of subjects have values *x_i_* in the vertex *x*. If a vertex has same values for all subjects, *V(x)* will be 0, indicating there is no variance of this vertex.

### Associations of network topography with development and executive function

We evaluated mass-univariate assocations between network topography and both development and executive function at two scales: total relative network representation and at each vertex. As an initial step, for each network, we summed the loading of all vertices to quantify the relative presence of this network on the cortex. Notably, as this was conducted in normalized template space, this measure was not impacted by individual differences in structural surface area. To model both linear and nonlinear developmental effects, we used generalized additive models (GAMs) with penalized splines (Wood, 2004). Importantly, the GAM estimates nonlinearities using restricted maximum likelihood (REML), penalizing nonlinearity in order to avoid over-fitting the data. We included sex and in-scanner head motion during scanning as model covariates. As we considered three functional runs, in-scanner motion was summarized as the grand mean of the mean relative RMS displacement of each functional run. To evaluate assocations with executive function, the executive function factor score was added as another model term with covariates as above (including a spline of age). Multiple comparisons were accounted for using the Bonferroni method.

To evaluate more complex topographic reconfiguration, we next evaluated each network at each vertex. For each network, we calculated associations between network loading and age for each vertex using GAMs while controlling sex and in-scanner head motion. Similarly, we calculated associations between network loadings and executive function while controlling sex, head motion, and a spline of age. Given the large number of multiple comparisons, for vertexwise analyses we used the False Discovery Rate (*Q* < 0.05).

### Prediction of brain maturity and executive function performance from spatial topography

Having tested if network topography was related to development and executive function in mass-univariate fashion, we next evaluated whether the overall multivariate spatial pattern of network topography encodes brain maturity or executive function. To address this question, we used ridge regression with nested two-fold cross validation (2F-CV, see **Fig. S3**) to test if multivariate network topography pattern could be used to identify an unseen individual participant’s age or executive function in an unbiased fashion. Accordingly, we combined the 17 network loading maps into a feature vector to represent the multivariate spatial pattern of network topography of each individual.

#### Ridge regression

A linear regression model was adopted to predict the brain maturity and executive function performance using the pattern of whole-brain spatial topography of parcellations. The linear model can be formalized as follows:

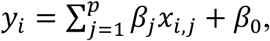

where *y_i_* is the age of the *i^th^* individual, *p* is the number of features, *x_i,j_* is the value of the *j^th^* feature of the *i^th^* subject, and *β_j_* is the regression coefficient.

To avoid over-fitting and to improve the prediction accuracy, we used ridge regression. Ridge regression uses an *L2* penalty during model fitting; we have previously shown often out-performs other methods for regression problems using high-dimensional imaging data and computationally more efficient than other methods (Cui and Gong, 2018; Hoerl and Kennard, 1970). The objective function is:

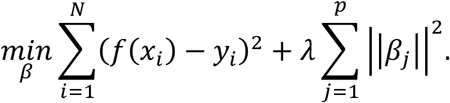

This technique shrinks the regression coefficients, resulting in better generalizability for predicting unseen samples. In this algorithm, a regularization parameter *λ* is used to control the trade-off between the prediction error of the training data and L2-norm regularization, i.e., a trade-off of penalties between the training error and model complexity. A large *λ* corresponds to a greater penalty on model complexity, and a small *λ* represents a greater penalty on training error.

#### Prediction framework

We applied a nested 2-fold cross validation (2F-CV), with outer 2F-CV estimating the generalizability of the model and the inner 2F-CV determining the optimal parameter *λ* for the ridge regression model (see **Figure S6** for schematic of the prediction framework).

#### Outer 2F-CV

In the outer 2F-CV, the data was divided into 2 subsets. Specifically, we sorted the subjects according to the outcome (i.e., age or executive performance) and then assigned the individuals with an odd rank to subset 1 and the individuals with an even rank to subset 2 (Cui and Gong, 2018; Cui et al., 2018). We intitally used subset 1 as the training set, with subset 2 used as the test set. Each feature was linearly scaled between zero and one across the training dataset, and the scaling parameters were also applied to scale the testing dataset (Cui and Gong, 2018; Erus et al., 2015). We applied an inner 2-fold cross validation (2F-CV) within training set to select the optimal *λ* parameter. Based on the optimal *λ*, we trained a model using all subjects in the training set, and then used that model to predict the outcome of all subjects in the testing set. Analogously, we used subset 2 as the training set and subset 1 as the test set, and repeated the above procedure. Across the testing subjects for each fold, the partial correlation and mean absolute error (MAE) between the predicted and actual outcome was used to quantify the prediction accuracy. In evaluation of the prediction of participant age, we controlled for sex and in-scanner head motion by calculating a partial correlation. Furthermore, we additionally controlled for participant age (in addition to sex and motion) when calculating the partial correlation between actual and predicted executive function. Here, we used the scikit-learn library to implement ridge regression (http://scikit-learn.org) (Pedregosa et al., 2011).

#### Inner 2F-CV

Within each loop of the outer 2F-CV, we applied inner 2F-CVs to determine the optimal *λ*. Specially, the training set for each loop of the outer 2F-CV was further partitioned into 2 subsets according to their rank of the outcome (i.e., age or executive performance), as in the outer loop (i.e., subjects with odd rank in subset 1 and subjects with even rank in subset 2). One subset was selected to train the model under a given *λ* in the range [2^−10^, 2^−9^, …, 2^4^, 2^5^] (i.e., 16 values in total) (Cui and Gong, 2018; Hsu et al., 2003), and the remaining subset was used to test the model. This procedure was repeated 2 times such that each subset was used once as the testing dataset, resulting in 2 inner 2F-CV loops in total. For each inner 2F-CV loop, the correlation *r* between the actual and predicted outcome and the mean absolute error (MAE) were calculated for each *λ*, and averaged over each fold. The sum of the mean correlation *r* and reciprocal of the mean MAE was defined as the inner prediction accuracy, and the *λ* with the highest inner prediction accuracy was chosen as the optimal *λ* (Cui and Gong, 2018; Cui et al., 2018). Of note, the mean correlation *r* and the reciprocal of the mean MAE cannot be summed directly, because the scales of the raw values of these two measures are quite different. Therefore, we normalized the mean correlation *r* and the reciprocal of the mean MAE across all values and then summed the resultant normalized values.

#### Significance of prediction performance

To evaluate if prediction performance (i.e., the partial correlation *r* and MAE) were significantly better than expected by chance, we performed a permutation test (Mourao-Miranda et al., 2005). Specifically, prediction procedure was re-applied 1,000 times. In each run, we permuted the outcome (i.e., age or executive function) across the training samples without replacement. The significance was determined by ranking the actual prediction accuracy versus the permuted distribution; the *p-*value of the partial correlation *r* was the proportion of permutations that showed a higher value than the actual value for the real data. Similarly, the *p-*value of the MAE was the proportion of permutations that showed a lower value than the actual value for the real data.

#### Interpreting model feature weights

The features with a nonzero regression coefficient/weight in the model trained using all subjects can be understood as contributing features for the prediction model (Cui and Gong, 2018; Mourao-Miranda et al., 2005). Absolute value of the weight quantified the contribution of the features to the model (Mourao-Miranda et al., 2005). To understand which network contributed the most to the prediction, we summed the absolute weight of all vertices in each network. Specifically, the vertices positively related to the outcome and that negatively related to the outcome were summed separately. We further tested if the veretices with small mean loadings, which were localized on the border of functional networks, contributed more to the prediction. We calculated the Pearson correlation between the absolute contribution weight and network loadings and used spin test to evaluate its significance. Finally, we tested if the spatial contribution of locations to the prediction was constrained by the variability of functional topography. As each vertex had 17 loading values (one for each network), we summed the absolute weight across all 17 networks to summarize the prediction weight of this vertex, which represents the importance of the vertex to the prediction, and then calculated the Pearson correlation between the summarized vertex weight and network variability across all vertices.

### Validation of multivariate prediction analysis

#### Randomly split 2F-CV

In the above prediction analysis, we split subjects into two halves according to the rank of the outcome (i.e., age or executive performance). To validate that our split was representative, we tested the prediction accuracy using repeated random 2F-CV. Specifically, we split the subjects randomly into two halves for both outer 2F-CV and inner 2F-CV, and calculated the mean partial correlation *r* and MAE across two folds. Because the split was random, we repeated this procedure 100 times and averaged the partial correlation (accounting for covariates) and MAE across the 100 times to determine the overall prediction accuracy. We used permutation testing to determine if the acquired prediction accuracy was significantly better than acquired by chance. Specifically, we repeated the random 2F-CV 1,000 times, but each time we permuted the outcome across the training data. Finally, we compared the actual mean partial correlation *r* and mean MAE to that of the null distrubtion.

#### Prediction by discrete network parcellations

Having demonstrated that continuously-weighted functional topography predicts age and executive performance, we next tested if the pattern of discrete network labels could predict age and executive performance. We extracted the whole-brain discrete network labels into a feature vector to represent the multivariate spatial pattern of network topography of each individual. Based on these features, we applied the above 2F-CV framework to predict age and executive performance using multivariate ridge regression. As network labels are categorical features, we first encoded each vertex feature as a one-hot numeric array (https://scikit-learn.org/stable/modules/generated/sklearn.preprocessing.OneHotEncoder.html), which was further used as input of the prediction analysis. This framework was applied to discrete networks from both NMF and MS-HBM.

#### Prediction of other cognitive measurements by network loadings

As a final step, we evaluated if the associations with network topography were specific to executive function. We used the same ridge regression framework and vertexwise loading maps to try to predict factor scores summarizing memory accuracy and social cognition accuracy.

### Visualization

Connectome Workbench (version: 1.3.2) provided by the human connectome project (https://www.humanconnectome.org/software/connectome-workbench; Marcus et al. (2013)) was used to visualize the brain surface.

### Data & code availability

The PNC data is publicly available in the Database of Genotypes and Phenotypes: accession number: phs000607.v3.p2; https://www.ncbi.nlm.nih.gov/projects/gap/cgi-bin/study.cgi?study_id=phs000607.v3.p2. All analysis code is available here: https://github.com/ZaixuCui/pncSingleFuncParcel, with detailed explanation in https://github.com/ZaixuCui/pncSingleFuncParcel/wiki.

## SUPPLEMENTARY FIGURES

**Supplementary Figure 1.**
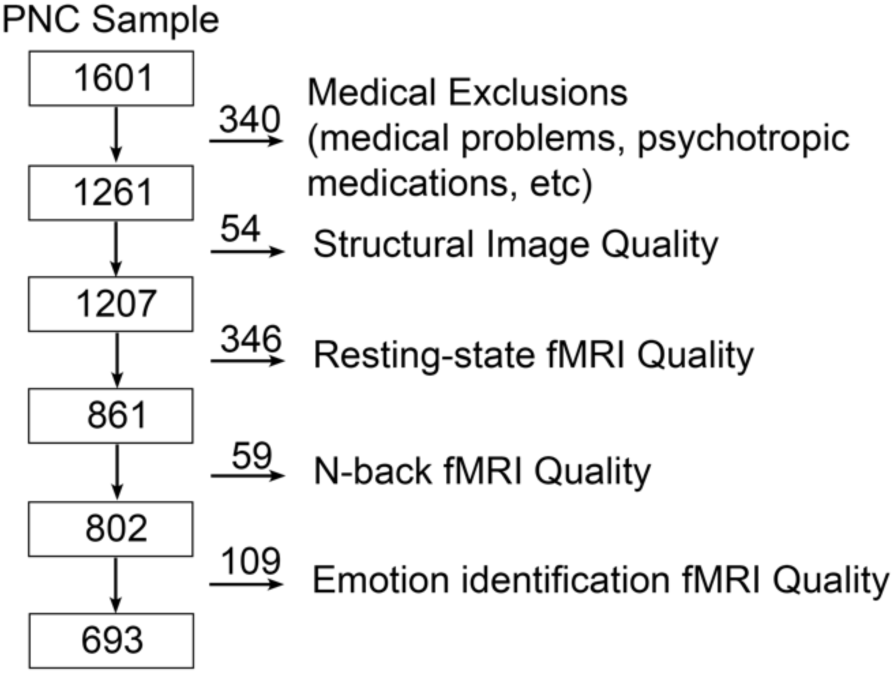
Sample construction. The Philadelphia Neurodevelopmental Cohort (PNC) included 1,601 participants who completed neuroimaging. Of these, 340 subjects were excluded owing to clinical factors, such as medical co-morbidity or use of psychotropic medication. Additionally, 568 subjects were excluded because of low quality or missing structural, resting-state, n-back, or emotion identification imaging data (details in Online Methods). The final sample consisted of the remaining 693 subjects.

**Supplementary Figure 2.**
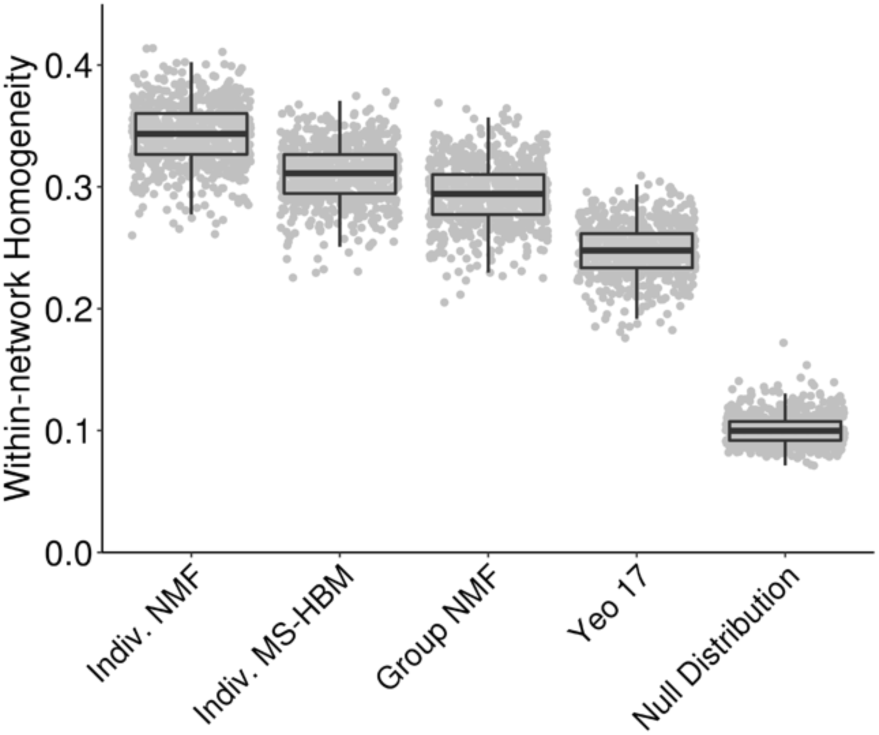
Individualized parcellation maximizes functional homogeneity within networks. The homogeneity of timeseries within each network is a common metric of parcellation quality. Both NMF and an alternative method for individualized network parcellation (MS-HBM) out-performed the group atlas from either NMF or the canonical 17 networks defined by (Yeo et al., 2011). All methods displayed greater within-network homogeneity than a null distribution of randomly rotated individualized networks that preserved the data’s spatial covariance structure.

**Supplementary Figure 3.**
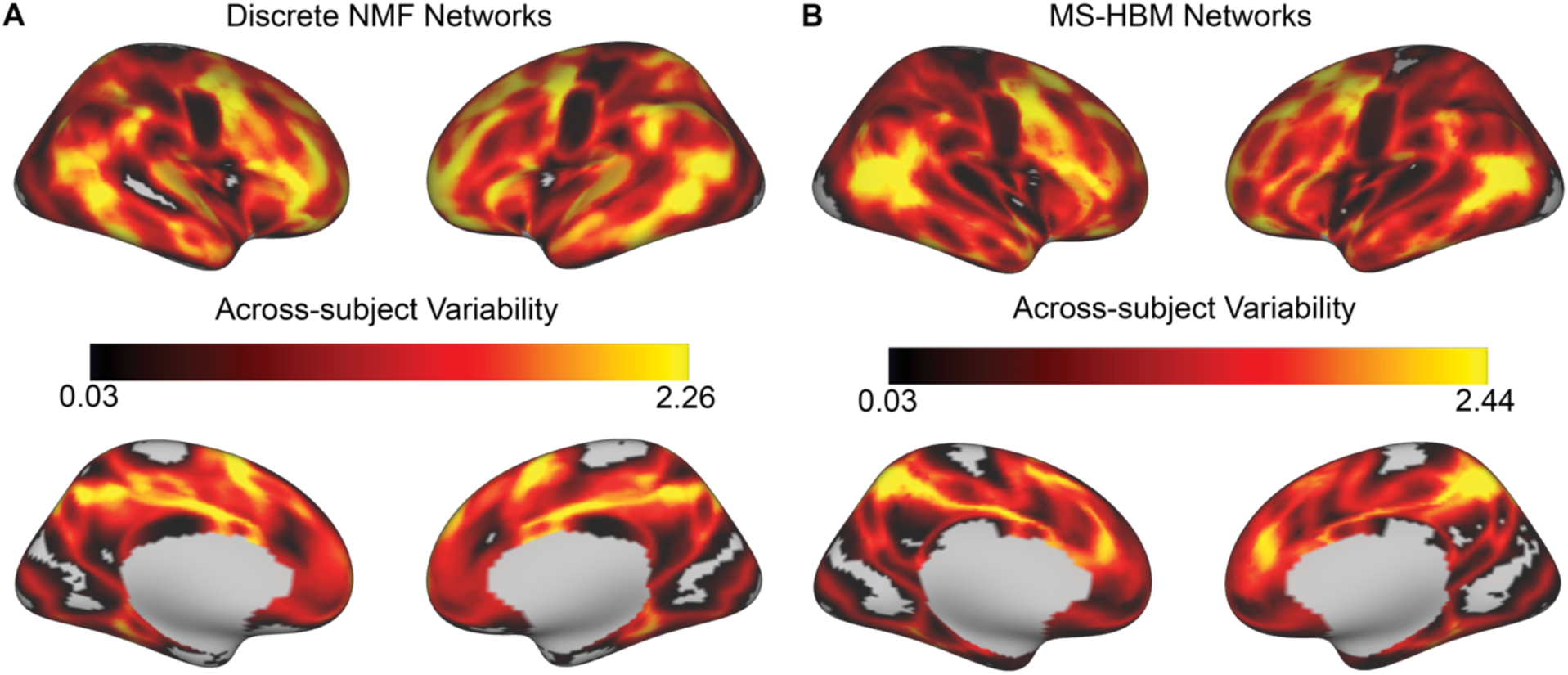
Across-subject variability of network topography of discrete network parcellations created by NMF (**A**) and MS-HBM (**B**).

**Supplementary Figure 4.**
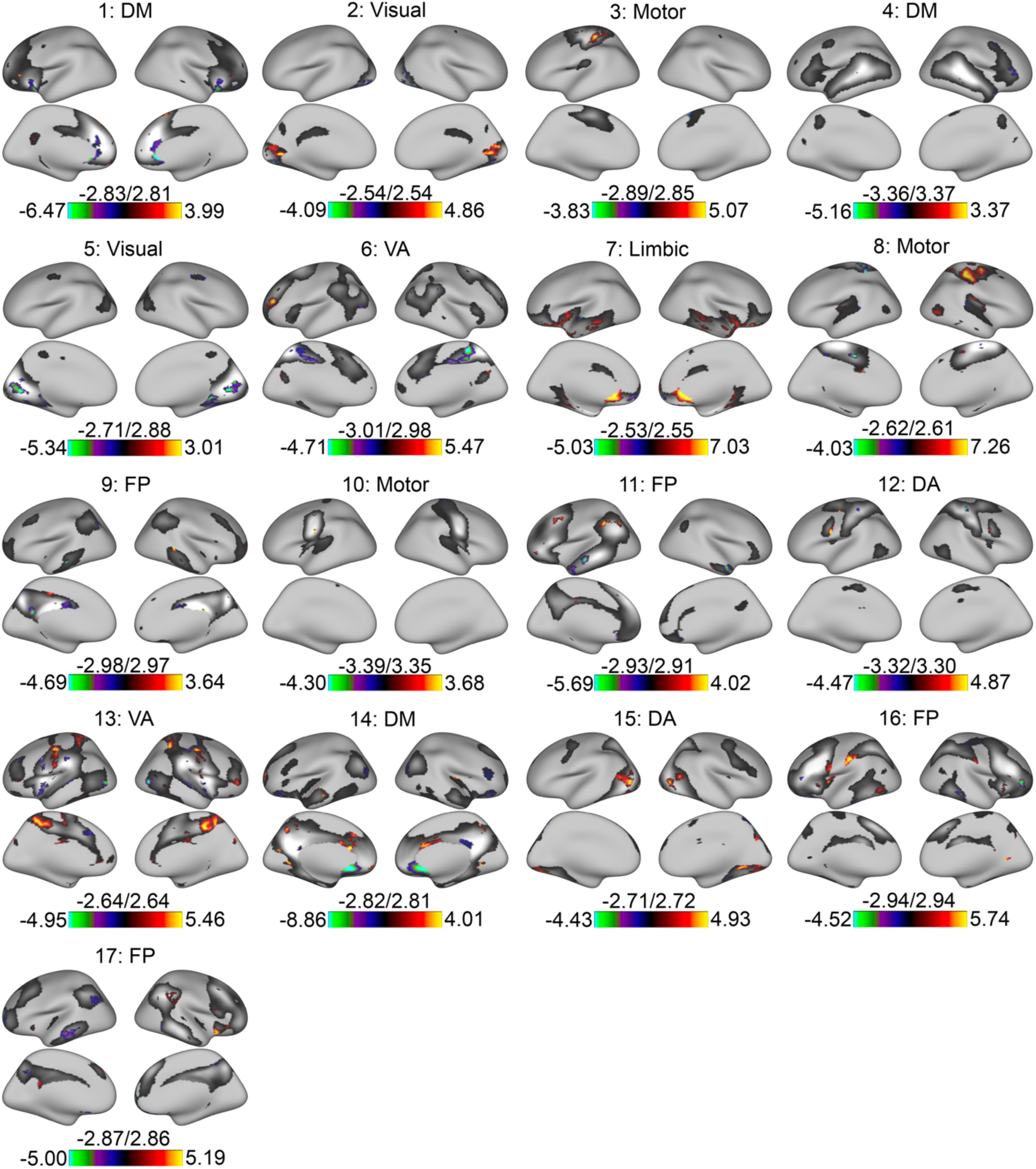
Vertex-wise associations with age for all networks (FDR *Q* < 0.05); underlay depicts group network atlas.

**Supplementary Figure 5.**
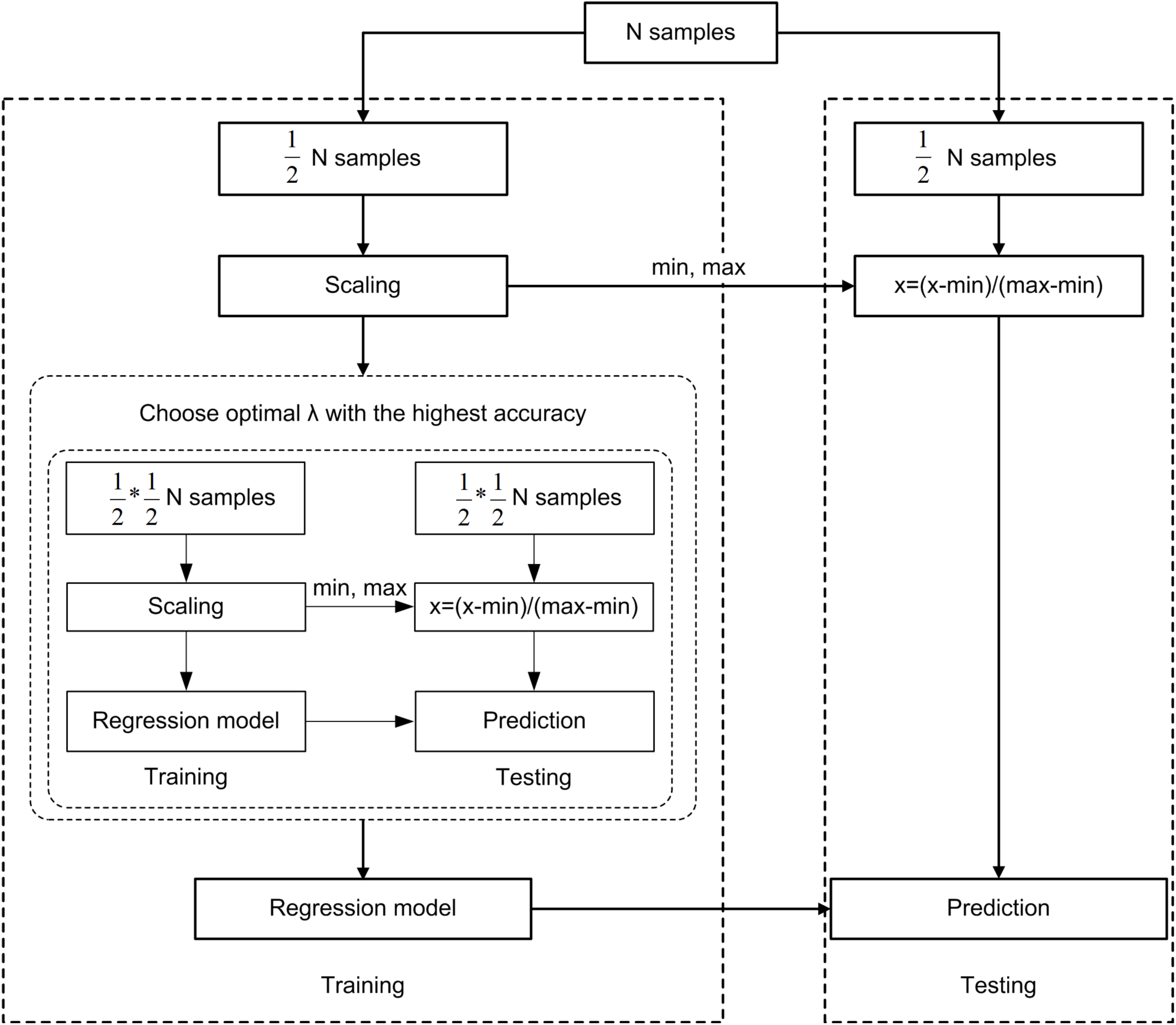
Schematic overview of one outer loop of the nested 2-fold cross-validation (2F-CV) prediction framework. All subjects were divided into two balanced halves according to the rank of the outcome (i.e., age or executive performance), with the first half used as a training set and the second half used as a testing set. Each feature was linearly scaled between zero and one across the training dataset; these scaling parameters were applied to the testing sample. An inner 2F-CV was applied within the training set to select the optimal *λ* parameter. Based on the optimal *λ*, we trained a model using all subjects in the training set, and then used that model to predict the outcome of all subjects in the testing set.

**Supplementary Figure 6.**
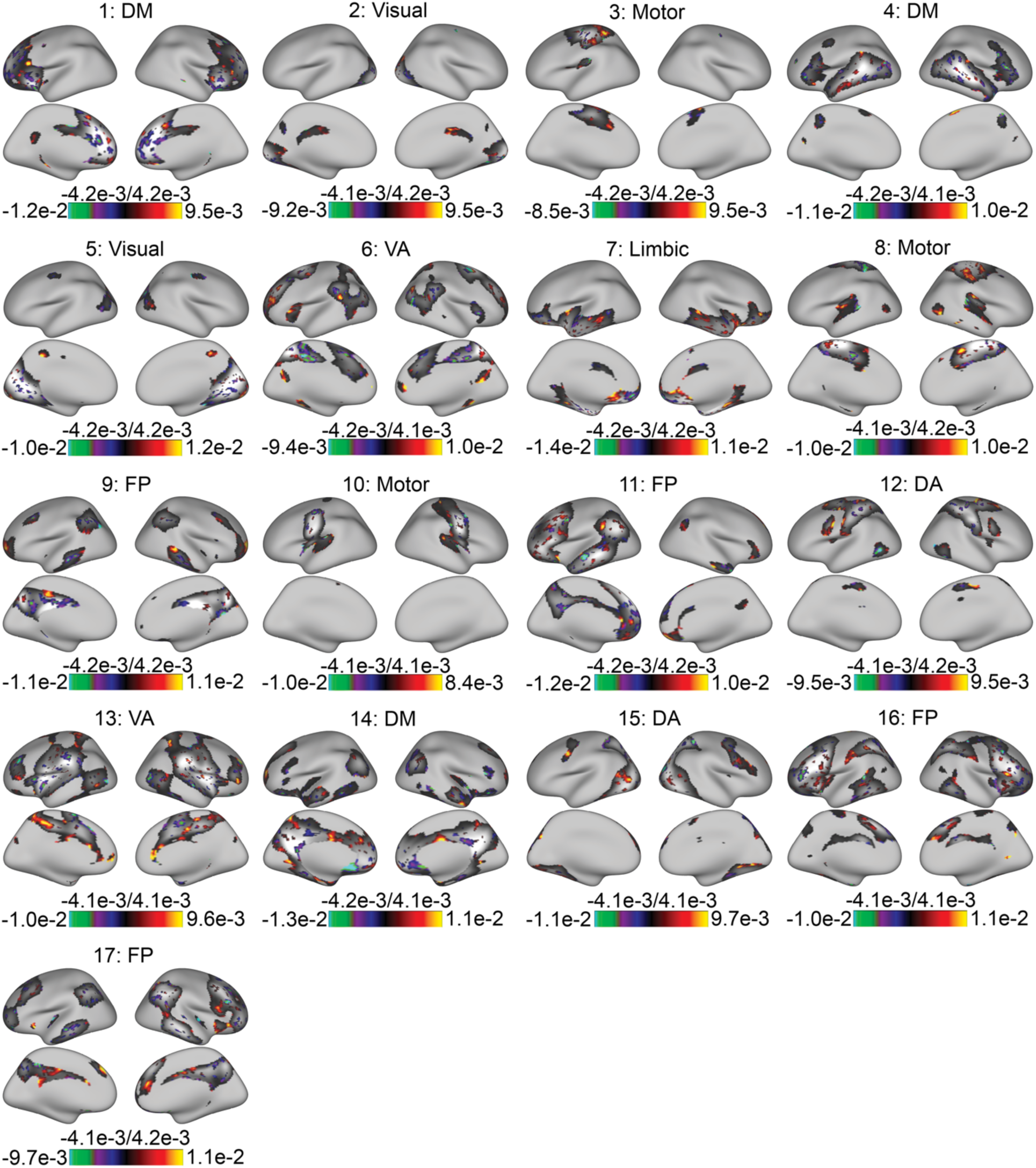
The vertices with the highest (first 25%) absolute regression weight in the multivariate prediction model of brain maturity; underlay depicts group network atlas.

**Supplementary Figure 7.**
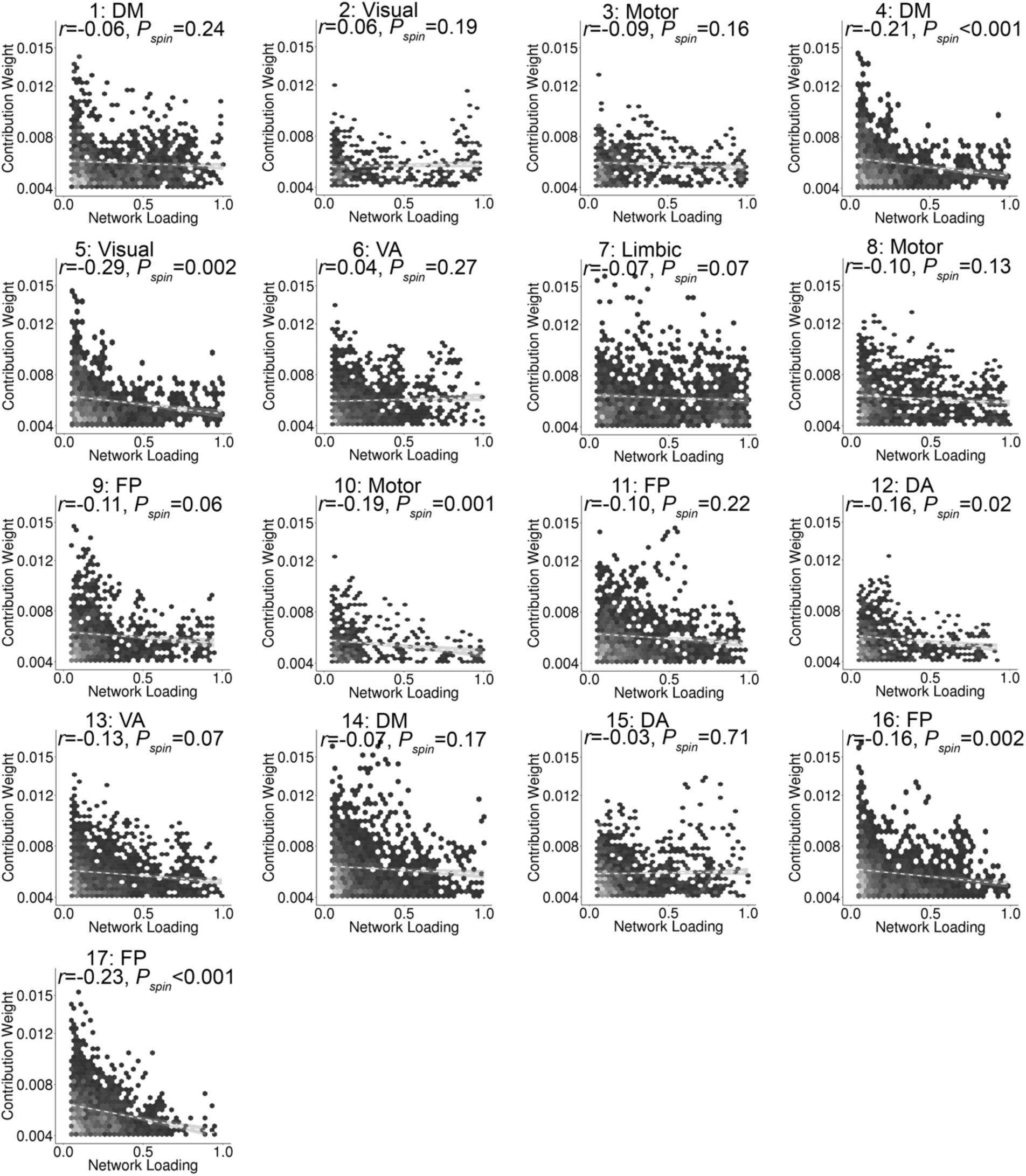
The absolute contribution weight (first 25%) was negatively correlated with group network loadings across vertices. Significance testing used the conservative spatial permutation procedure (spin test).

**Supplementary Figure 8.**
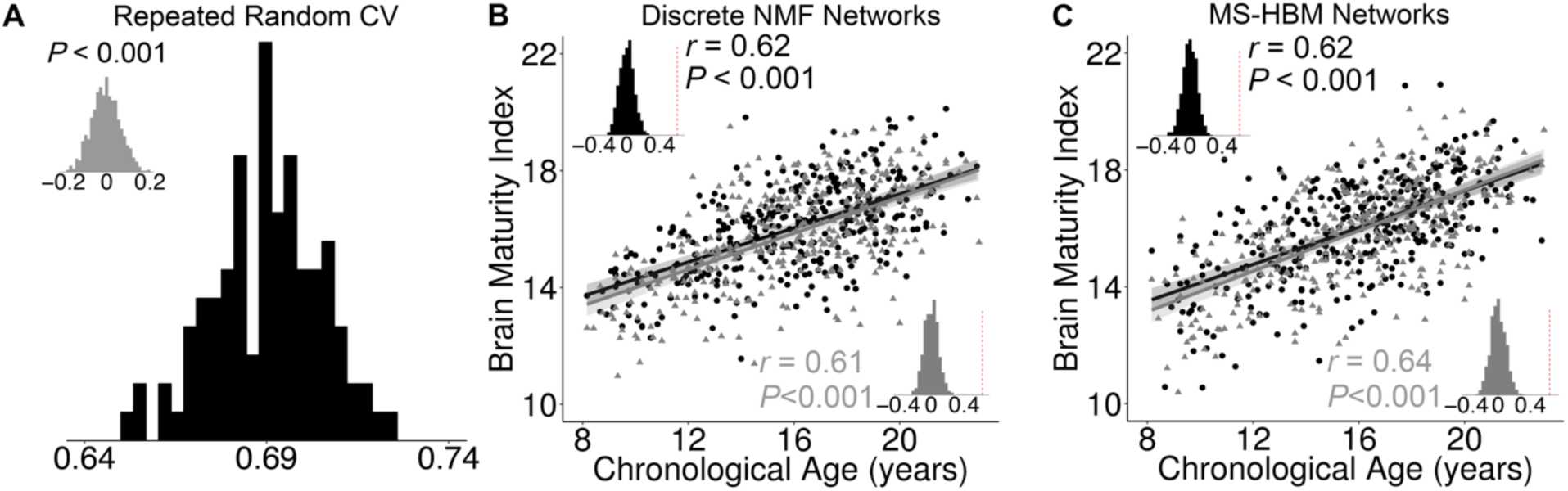
Supplementary analyses for brain maturity prediction using multivariate pattern of functional topography. (**A**) Evaluated by repeated random 2-fold cross-validation (2F-CV), functional topography of network loadings predict an unseen individual’s brain maturity significantly higher than by chance. Black histogram is the distribution of prediction accuracy of repeated random 2F-CV, and the inset histogram displays the distribution of prediction accuracy from a permutation test. There was no overlap between these distributions. (**B**) Functional topography of discrete NMF parcellations predict brain maturity. (**C**) Functional topography of discrete MS-HBM parcellations predict brain maturity.

**Supplementary Figure 9.**
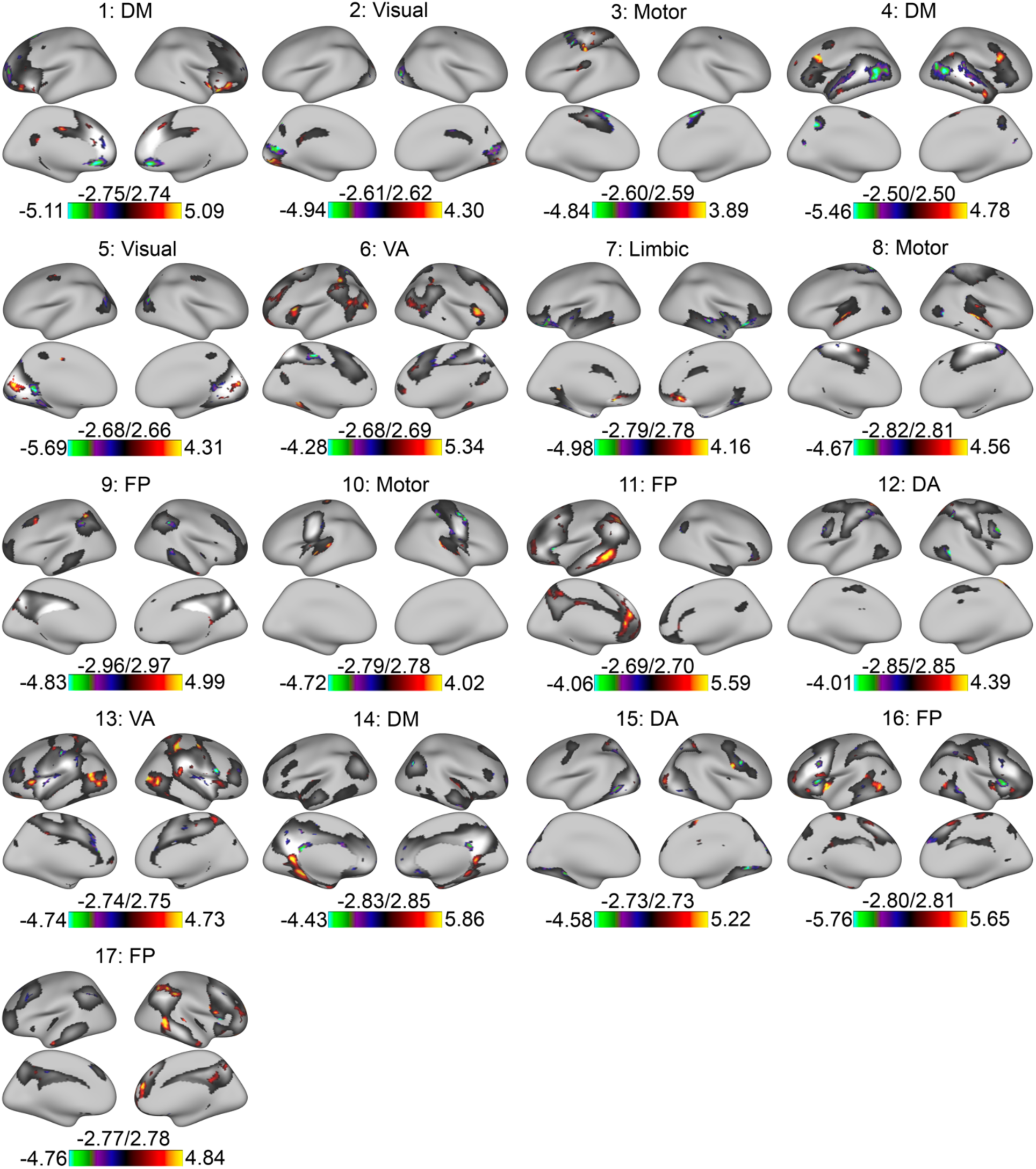
Vertex-wise associations with executive performance for all networks (FDR *Q* < 0.05); underlay depicts group network atlas.

**Supplementary Figure 10.**
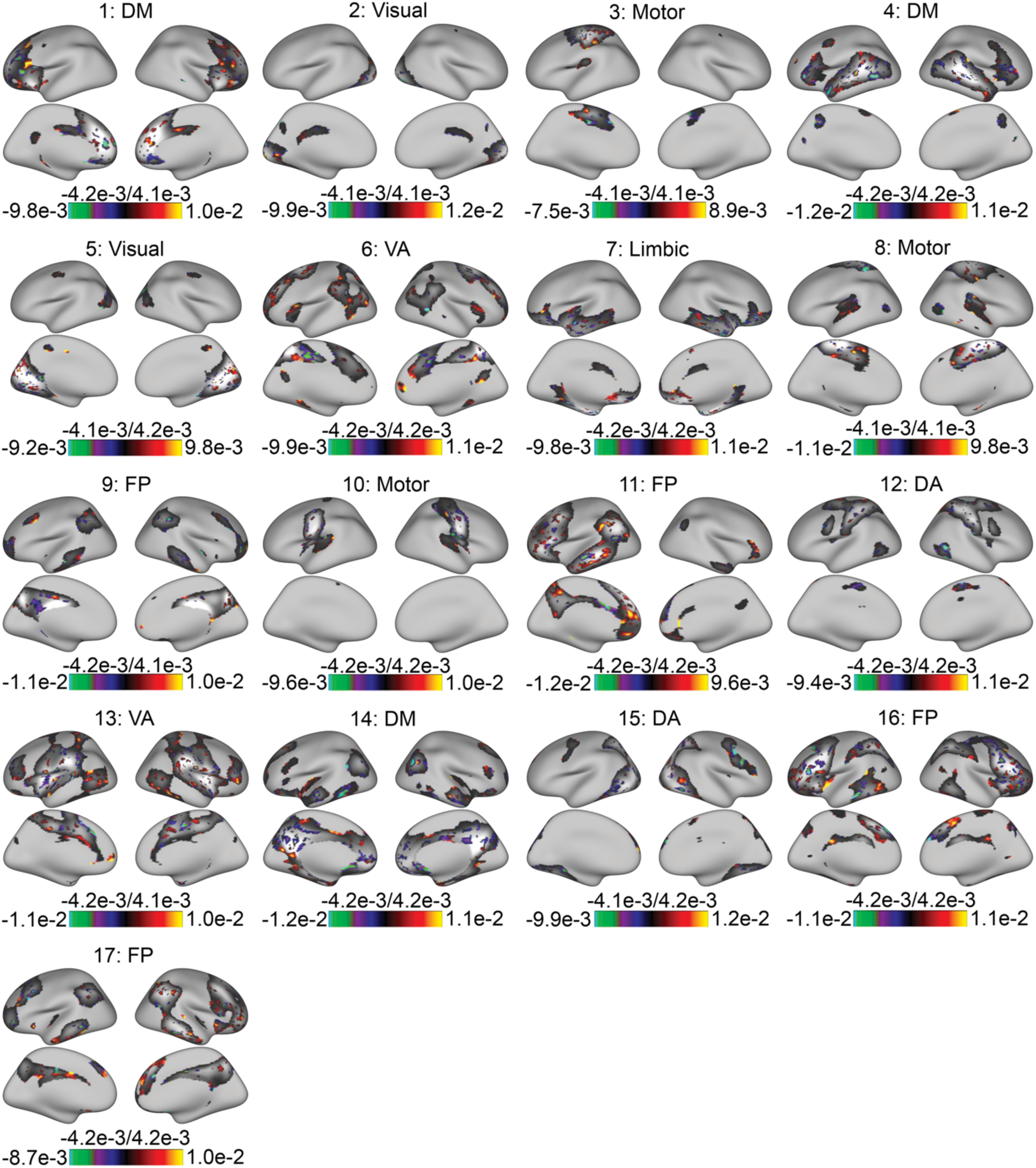
The vertices with the highest (top 25%) absolute regression weight in the multivariate prediction model of executive performance; underlay depicts group network atlas.

**Supplementary Figure 11.**
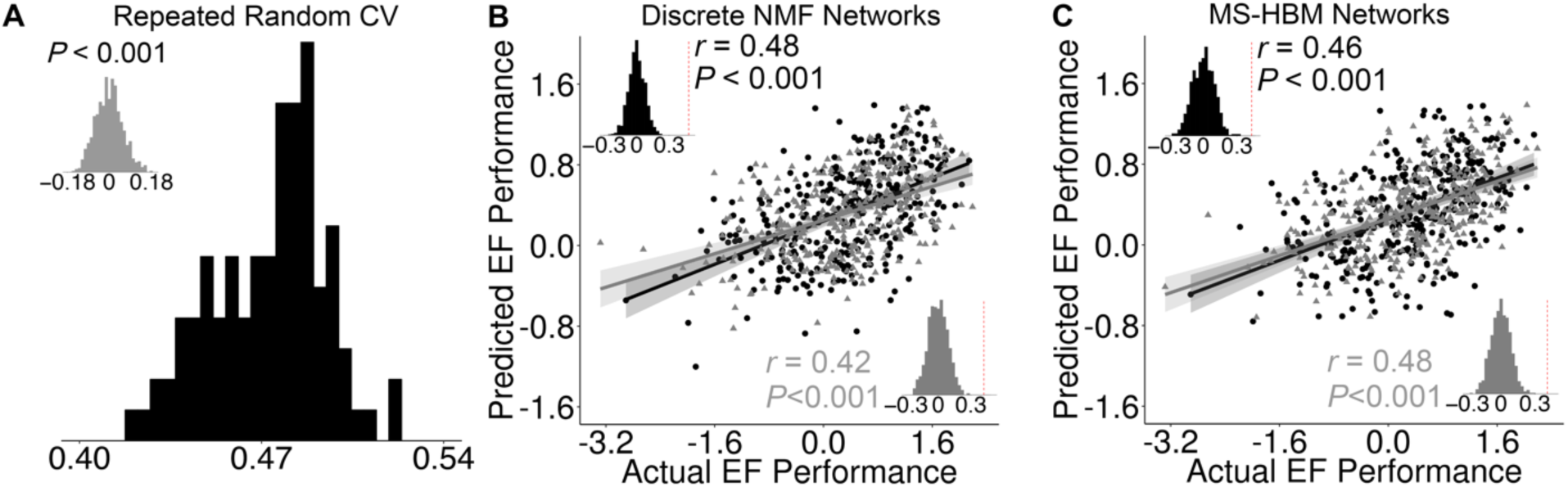
Supplementary analyses of executive function prediction using multivariate patterns of functional topography. (**A**) Evaluated by repeated random 2-fold cross-validation (2F-CV), functional topography of network loadings predict an unseen individual’s executive performance significantly higher than by chance. Black histogram is the distribution of prediction accuracy of repeated random 2F-CV, and the inset histogram displays the distribution of prediction accuracy from a permutation test. There was no overlap in these distributions. (**B**) Functional topography of discrete NMF parcellations predicts unseen individuals’ executive performance. (**C**) Functional topography of discrete MS-HBM parcellations predicts unseen individuals’ executive performance.

